# Dual G9A and EZH2 inhibition stimulates an anti-tumour immune response in ovarian high-grade serous carcinoma

**DOI:** 10.1101/2021.05.09.443282

**Authors:** Pavlina Spiliopoulou, Sarah Spear, Hasan Mirza, Ian Garner, Lynn McGarry, Fabio Grundland-Freile, Zhao Cheng, Darren P. Ennis, Sophie McNamara, Marina Natoli, Susan Mason, Karen Blyth, Peter D. Adams, Patricia Roxburgh, Matthew J. Fuchter, Bob Brown, Iain A. McNeish

## Abstract

Ovarian high-grade serous carcinoma (HGSC) prognosis correlates directly with presence of intratumoral lymphocytes. However, cancer immunotherapy has yet to achieve meaningful survival benefit in patients with HGSC. Epigenetic silencing of immunostimulatory genes is implicated in immune evasion in HGSC and re-expression of these genes could promote tumour immune clearance. We discovered that simultaneous inhibition of the histone methyltransferases G9A and EZH2 activates the CXCL10-CXCR3 axis and increases homing of intratumoral effector lymphocytes and natural killer cells whilst suppressing tumour-promoting FoxP3^+^ CD4 T cells. The dual G9A/EZH2 inhibitor HKMTI-1-005 induced chromatin changes that resulted in the transcriptional activation of immunostimulatory gene networks, including the re-expression of elements of the ERV-K endogenous retroviral family. Importantly, treatment with HKMTI-1-005 improved the survival of mice bearing *Trp53*^*-/-*^null ID8 ovarian tumours and resulted in tumour burden reduction.

These results indicate that inhibiting G9A and EZH2 in ovarian cancer alters the immune microenvironment and reduces tumour growth and therefore positions dual inhibition of G9A/EZH2 as a strategy for clinical development.

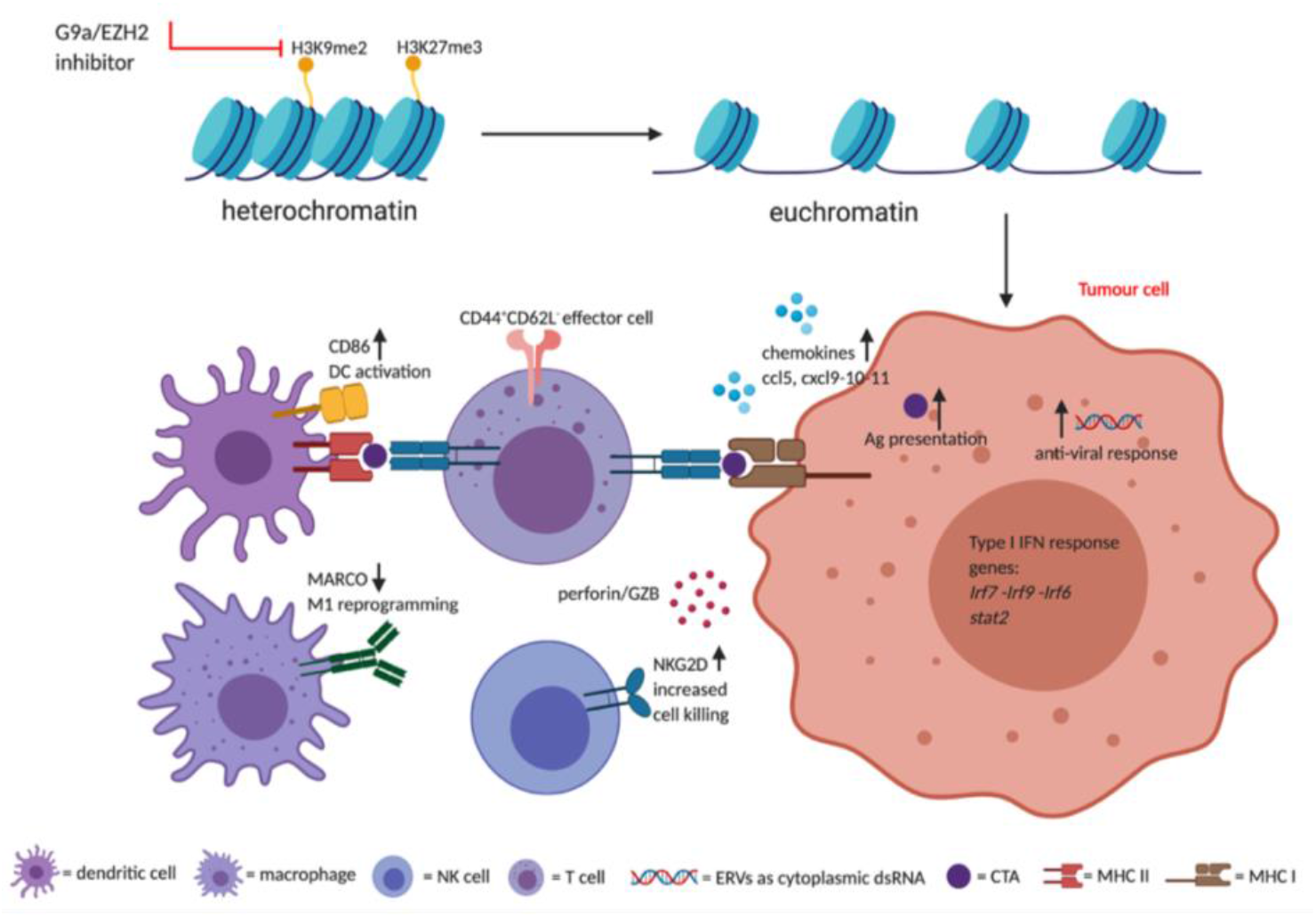

## Introduction

Despite ample evidence that the prognosis of patients with advanced ovarian high-grade serous carcinoma (HGSC) is strongly influenced by the immune microenvironment (1), current immunotherapies have failed to produce a meaningful survival benefit for patients (2). HGSC cells can evade immune responses by altering their epigenome, and targeting ovarian cancer epigenetics can reactivate cancer testis antigens (3-5), induce viral mimicry (6, 7) and alter the tumour immune microenvironment and immune cell function (8, 9). DNA methylation and histone deacetylation are two mechanisms that play a role in cancer immune evasion (10, 11), and, although inhibitors of both DNA methylation and histone deacetylation are currently used in some haematological malignancies, their use in solid malignancies has been limited due to toxicity and limited efficacy (12, 13). More recently, histone methylation mediated by both G9A and EZH2 has been identified as an important pathway that influences the immune system in ovarian cancer and melanoma, as well as hepatocellular, multiple myeloma and bladder carcinomas (8, 14-18).

Increased levels of chemokines the CXCL9, CXCL10, CXCL11 and CCL5 are all associated with an immune-reactive ovarian cancer (OC) microenvironment and improved patient prognosis (19). CXCL9, CXCL10 and CXCL11 are interferon-inducible and bind to the receptor CXCR3. Genomic analysis of 489 patient samples by The Cancer Genomic Atlas (TCGA) Research Network identified a defined subgroup of OC patients with an activated CXCR3/CXCL9-11 pathway (20) and, critically, when these chemokines are present at high concentrations in tumours, patients achieve a longer disease-free interval and overall survival (21, 22). The primary role of these IFN-γ-inducible chemokines is trafficking of activated CD8^+^, CD4^+^ T cells and natural killer (NK) cells. In preclinical models of OC, increased expression of CXCL10 can reduce tumour burden and ascites accumulation (23). CCL5 is also associated with T cell infiltration and tumour control in oesophageal and hepatocellular carcinomas (24, 25). Coukos et al. recently showed that CCL5^hi^CXCL9^hi^ ovarian tumours are immunoreactive and responsive to immune checkpoint blockade, with tumour-derived CCL5 driving expression of CXCL9 from intratumoral immune cells, such as antigen-presenting cells, which in turn support T cell engraftment in the tumour (26). We reasoned that pharmacological approaches to activate the CXCR3/CXCL9-11 pathway might be of therapeutic benefit in OC.

Using a medium-throughput screening library of epigenetic compounds, we sought to discover novel epigenetic mechanisms that can augment immune responses in HGSC. We discovered that dual inhibition of G9A and EZH2 histone lysine methyltransferases induces potent release of lymphocyte chemotactic chemokines, including CXCL9, CXCL10, CXCL11 and CCL5, confirming these results in a panel of human cell lines and primary patient samples. We also showed that the dual G9A/EZH2 inhibitor HKMTI-1-005 (27) powerfully modified accessible chromatin in a syngeneic HGSC model, accompanied by transcriptional upregulation of immune pathways and, critically, substantial modulation of the tumour immune microenvironment. Importantly, we describe how G9A/EZH2 inhibition generated a significant influx of effector CD8^+^ T cells, NK cells, activated conventional type 1 dendritic cells (cDC1) whilst depleting tumours of CD4^+^ T regulatory cells. We observed a significantly extended survival of mice treated with HKMTI-1-005, indicating that G9A/EZH2 inhibition may provide a useful tool to overcome the poor immune reaction to OC in patients.

## Methods

### In vivo syngeneic mouse model of ovarian cancer

All experiments performed in mice were approved by the Animal Welfare & Ethics Review Body (AWERB) at the University of Glasgow and Imperial College London. Experiments were performed under the project licence numbers 70/8645 at the Cancer Research UK Beatson Institute and project licences 70/7997 and PA780D61A at Imperial College London. All experiments conformed to UK Home Office regulations under the Animals (Scientific Procedures) Act 1986, including Amendment Regulations 2012. The study was compliant with all relevant ethical regulations regarding animal research. Humane endpoints included: weight loss of 20% or more, ascites equivalent to full term pregnancy, reduced/slow activity, pale feet and visible symptoms of distress such as hunching, piloerection, closed eyes and isolation from cage mates.

For both *in vitro* and *in vivo* experiments, we utilised the ID8 syngeneic murine model (28) with bi-allelic *Trp53* deletions that we previously described (29). *In vivo*, 5×10^6^ *Trp53*^*-/-*^ID8 cells/mouse were injected intraperitoneally (IP) in 6-week-old C57BL/6J female mice (Charles River, UK). At defined endpoint, ascites, intra-abdominal tumours (formed in omentum and porta hepatis) and spleens were collected (Figure S1). When no ascites was present, peritoneal cells were collected by lavage with 5 mL phosphate-buffered saline (PBS). HKMTI-1-005 was given as twice daily IP injections of 20 mg/kg. HKMTI-1-005 (30) was dissolved in DMSO for long term storage and reconstituted in 1% Tween/0.9% NaCl vehicle, just prior to injection. Mice were randomly assigned to a 2-week treatment with HKMTI-1-005 or vehicle alone (control) starting on day 21 following IP cell inoculation. The investigators deciding on endpoint were blinded to the treatment administered.

### Drug library screening

The drug library of epigenetic compounds was provided by Structural Genomic Consortium (Oxford) (Table S1). 2×10^4^ *Trp53*^*-/-*^ID8 cells/per well were seeded in 364-well black polypropylene, flat-bottom plates on day -1. On day 0, the drug library was added, along with 1 ng/mL of mouse IFNγ (ThermoFisher #PMC4031). After 72 hours, supernatant was transferred onto CXCL10 enzyme-linked immunosorbent assay (ELISA) plates (R&D Systems DY466) using the JANUS G3 MDT automated workstation (Perkin Elmer) for downstream analysis (R&D Systems, DY466). Cell viability was measured by 4′,6-diamidino-2-phenylindole (DAPI) staining.

### Gene expression assays

RNA was extracted from cells with the Qiagen RNeasy protocol (Cat. No. 74004). Quality control and quantification were performed using the NanoDrop 2000 spectrophotometer (Thermo Scientific, Wilmington, DE, USA). RNA was aliquoted and stored at -80°C. cDNA synthesis was performed using High-capacity cDNA reverse transcription kit from ThermoFisher (4368814) and iTaq/universal probes mastermix (Bio-Rad, 1725131) was used for single-gene RT-qPCR reactions (Table S1). Chemokine expression was quantified using RT^2^ Profiler PCR array for mouse chemokines/cytokines (Qiagen PAMM-150ZA, 330231) with data analysis performed using the Qiagen online tool PCR Array data analysis Web portal (https://www.qiagen.com/gb/resources/resourcedetail?id=20762fd2-8d75-4dbe-9f90-0b1bf8a7746b&lang=en). The chemokines tested, quality control and normalisation analysis are found in Tables S3-S5.

### Human ascites

Ascites from patients with HGSC was collected under the approval of Imperial College Healthcare NHS Trust Tissue Bank (ICHTB HTA licence: 12275). After sterile collection, spheroids were captured on a 40 μm membrane and placed into T75 ultra-low attachment flasks (ULA, Corning 3814) and cultured in advanced DMEM/F12 medium (Life Technologies, 12634010), supplemented with 10% autologous ascites, 10 mM HEPES, 1x N-2 supplement (ThermoFischer, 17502048), 1x serum-free B-27 supplement (ThermoFischer, 17504044), 100 U/mL penicillin plus 100 μg/mL streptomycin (penicillin/streptomycin Thermofisher, 15140-122), and 2 mM L-glutamine (Thermofisher, 25030-081). Spheroids were allowed to grow for up to 72-96h after which they were dissociated and treated as a monolayer. Patient details are given in Table S6.

### Flow cytometry

Fresh tumour deposits from the omentum and the porta hepatis of mice bearing *Trp53*^*-/-*^ID8 tumours were harvested. Tumour digestion was performed as previously described (31) modified with the use of collagenase (Sigma, C7657) and dispase (Gibco, 17105041). Tumour cells (20×10^6^/mL) were plated in 96 well V-bottomed plates followed by FcR II/III block [BD Biosciences, 553142, diluted 1:200 in FACS (0.5% FBS, 2 mM EDTA in PBS) buffer]. Antibody details are given in Table S7. Cells were fixed with 2% neutral-buffered formalin diluted in FACS buffer following addition of Zombie Yellow fixable viability dye (BioLegend #423103, 1:200 in PBS).

For intracellular assessment of T cell activity, 20×10^6^/mL tumour cells were plated in clear untreated U-bottom plates (SLS, 3879) after tumour digestion. After stimulation (PMA and ionomycin, eBioscience, 00-4970, 2 μl/mL, 1h), protein transport inhibitor cocktail (eBioscience, 00-4980, 2 μl/mL) was added. After 4 hours, cells were transferred to a V-bottom plate and stained for the membrane markers, viability dye Zombie Yellow and fixed followed by intracellular staining using the intracellular staining permeabilization buffer (BioLegend, 421002). Samples were analysed on a 3-LASER Cytek^®^ Aurora, (Cytek^®^ Biosciences) cytometer and the software FlowJo™ 10.7.1. Only samples that reached a threshold of 200 events per sample were included in the quantitative analysis. The geometric mean fluorescence intensity (MFI) was calculated by subtracting by an average (minus FMO) fluorescence value from pooled samples from each individual test sample.

### RNA and ATAC sequencing

Frozen mouse tumours (≤10mg) were homogenised in a Precellys homogeniser using ceramic beads at 2,000 x g for 2 pulses of 30 seconds. RNA was extracted from the lysate with the Qiagen RNeasy protocol (Cat. No. 74004). RNA with an RNA Integrity number (RIN) of >7 as measured in an Agilent 2200 TapeStation was used for downstream sequencing analysis. Following ribosomal RNA depletion (NEBNext, E6350) from 250 ng total RNA, sequencing libraries were constructed using the NEBNext Ultra II Directional Library prep kit for Illumina (E7760S). Following QC (Agilent D5000 Screen Tape System) and quantification [Qubit dsDNA high sensitivity assay kit (Thermofisher, Q32854)], samples were sequenced (Nova6000 SP flow cell (Illumina) 50 bp PE, target 50 million read pairs per sample). For the assay of transposase-accessible chromatin using sequencing (ATAC-seq), the published protocol Omni-ATAC (32) was optimised for *Trp53*^*-/-*^ID8 tumours. 20 mg tumour deposits were homogenised with a glass Dounce homogeniser. The lysate was then mixed in an iodixanol concentration gradient (25%-29%-35% concentration gradient, iodixanol Sigma-Aldrich, <u>D1556</u>) and centrifuged in a swinging bucket at 4,000 x g for 20 minutes. After centrifugation, 20,000 nuclei per sample were harvested from the nuclear band and treated with 100 nM hyperactive transposase enzyme (Nextera Tagment DNA enzyme I #15027916) for 30 minutes at 37°C, shaking at 400 rpm, in an Eppendorf Thermomixer comfort incubator. The purified, transposed DNA was amplified using customised primers as previously published (33). After quality control (Agilent D5000 Screen Tape System), the amplified library was sequenced (Nova6000 S1 flow cell (Illumina), 50 bp PE).

Raw sequencing reads were aligned to mouse genome version GRCm38.p4 (mm10) using the STAR aligner with default parameters (34). Raw counts were generated using the Rsubread package (35) and Differentially Expressed Genes (DEGs) were identified using the DESeq package (36). All analyses, statistical tests, and plots were generated in R version 3.3.3 unless specified otherwise. MultiQC was used to collate data across different programs (37). For functional annotation of DEGs, we used the Database for Annotation, Visualization and Integrated Discovery (DAVID) online Functional Annotation Tool (38) with access to Gene Ontology (GO) (39), and KEGG (40) databases. For analysis of endogenous retroviruses (ERVs), a mm10 annotation for mouse endogenous viral elements was obtained from the gEVE database (41). During alignment, only primary alignments were taken into account, a method adapted by Haase et al (42). For ATAC-seq methodology, the MACS2 tool was used to call peaks on all individual control and treatment samples (43).

### Statistics

Statistical analyses were performed in GraphPad (PRISM 9.0.0 (86)). For mean comparisons between 2 groups, *t*-test was used for populations with normal distribution and Mann-Whitney test for non-parametric distribution. One-way ANOVA was used for comparison between more than 2 groups. Wilcoxon matched-pair test was used to compare median values for patient samples. Log-rank test was used to compare differences in survival. When indicated, ROUT (Q=1%) method was used to identify outliers.

## Results

### Combined G9A/EZH2 inhibition upregulates chemotactic chemokines in vitro

We initially screened 38 epigenetic drugs for the production of CXCL10 by IFN-γ stimulated *Trp53*^*-/-*^ID8 (29) cells using ELISA. IFN-γ induces *Cxcl10* transcription. We wanted to discover chromatin modifying drugs that could potentially enhance this induction by IFN-γ. A single concentration screen of all 38 drugs highlighted candidate compounds that increased CXCL10 production (**Fig 1A**). A two-dose re-screen (**Fig 1B**) indicated that UNC0642, an inhibitor of G9A (EHMT2) and G9A-like protein (GLP) (44), significantly upregulated CXCL10 production compared to IFN-γ stimulation alone (mean fold change 16.1 ± 7.4 *vs* 1.8 ± 0.3, p<0.0001) (**Fig 1B/Fig 1C**), at doses that were not cytotoxic (dose response curves, **Fig S2**).

**Figure 1:**
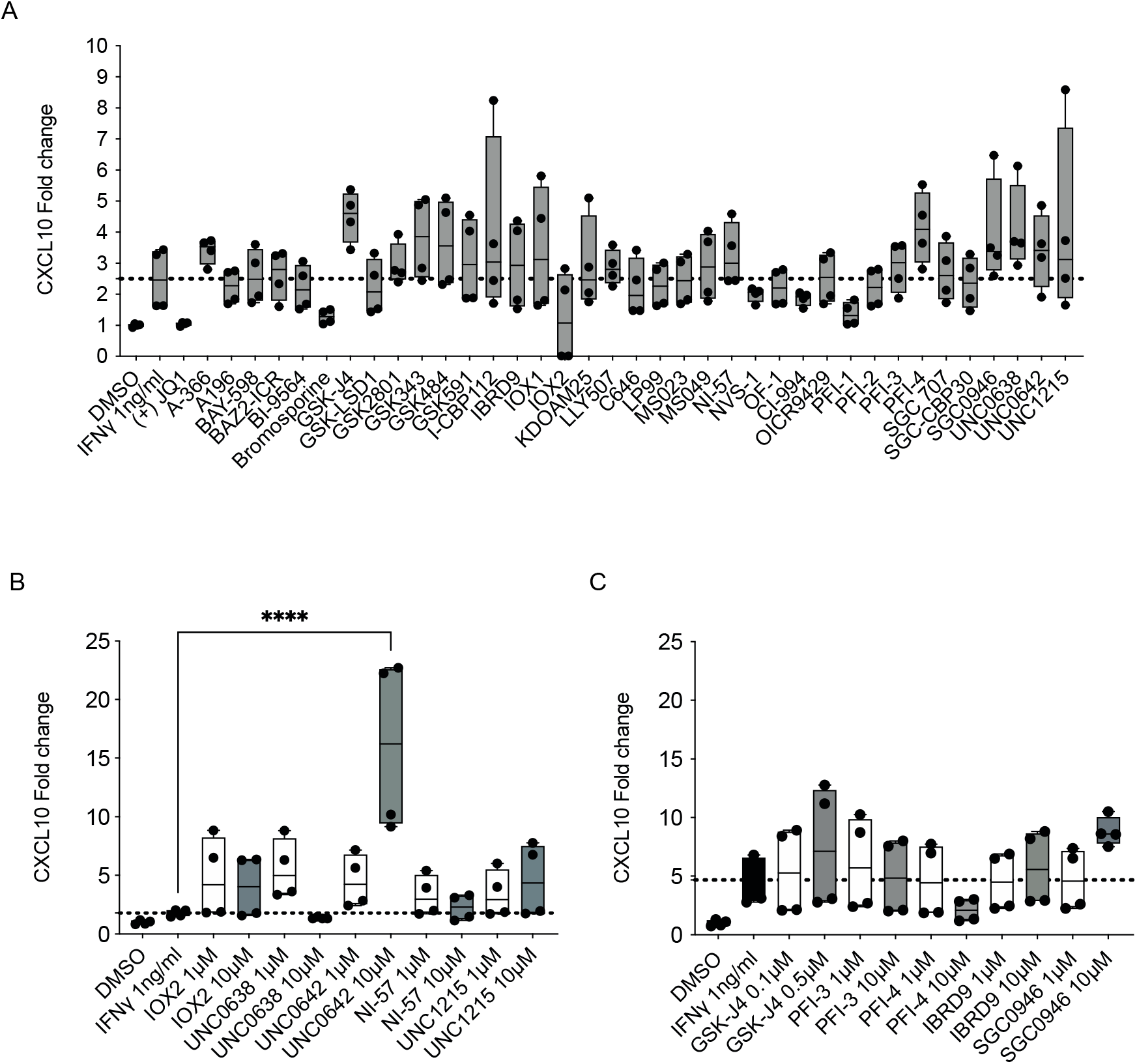
G9A inhibition upregulates CXCL10 in an ovarian cancer model. A. 2×10^3^ *Trp53*^*-/-*^ID8 cells in 384 well plates were treated with the SGC drug library (all drugs were at a final concentration of 1 μM apart from GSK-J4 -0.2 μM) with 1ng/ml of mouse IFN-γ. CXCL10 ELISA was performed on day 4. Box and whiskers show all values obtained from four technical replicates. Mean values were compared to IFN-γ stimulation alone, using one-way ANOVA with Dunnett’s multiple comparisons test. B and C. CXCL10 protein fold change following treatment with 10 selected SGC library drugs (1 μM and 10 μM apart from GSK-J4 - 0.1 μM and 0.5 μM). B and C show data from separate experiments. Box and whiskers show all values obtained from four technical replicates. One-way ANOVA with Dunnett’s multiple comparison test was used to compare all mean values to IFN-γ alone; statistically non-significant results are not shown. ****p <0.0001

As G9A cooperates closely with Enhancer of zeste homolog 2 (EZH2) in some contexts (45), we combined UNC0642 with an EZH2 inhibitor, UNC1999; this combination induced a greater increase of both *Cxcl10* mRNA and CXCL10 protein than either drug alone (mRNA mean fold change 109.4 ± 25.0 *vs* 12.3 ± 0.67 *vs* 12.49 ± 3.1, p<0.0001; protein mean fold change 2.23 ± 0.07 *vs* 1.9 ± 0.01 *vs* 1.4 ± 0.01, p< 0.0001, **Fig 2A, 2B**). As simultaneous inhibition of G9A and EZH2 achieved the most potent CXCL10 induction, we then evaluated the dual G9A/EZH2 inhibitor HKMTI-1-005, the first described dual inhibitor of its class (27, 30). HKMTI-1-005, in contrast to other known EZH2 inhibitors, has a peptide substrate competitive mechanism. At doses that reduced repressive histone marks (**Fig S3**), HKMTI-1-005 induced stronger upregulation of *Cxcl10* than either methyltransferase inhibitor alone (mean fold change 159.6 ± 12.5 *vs* 12.3 ± 0.67 *vs* 12.49 ± 3.1, p<0.0001, **Fig 2A**), as well as combination treatment with the 2 single inhibitors (mean fold change 159.6 ± 12.5 *vs* 109.4 ± 25.0, p=0.001, **Fig 2A**). HKMTI-1-005 treatment also resulted in higher CXCL10 protein production than the individual inhibitors given alone (mean fold change 3.1 ± 0.03 *vs* 1.9 ± 0.01 *vs* 1.4 ± 0.01, p<0.0001) and in combination (mean fold change 3.1 ± 0.03 *vs* 2.2 ± 0.1, p<0.0001, **Fig 2B**).

**Figure 2:**
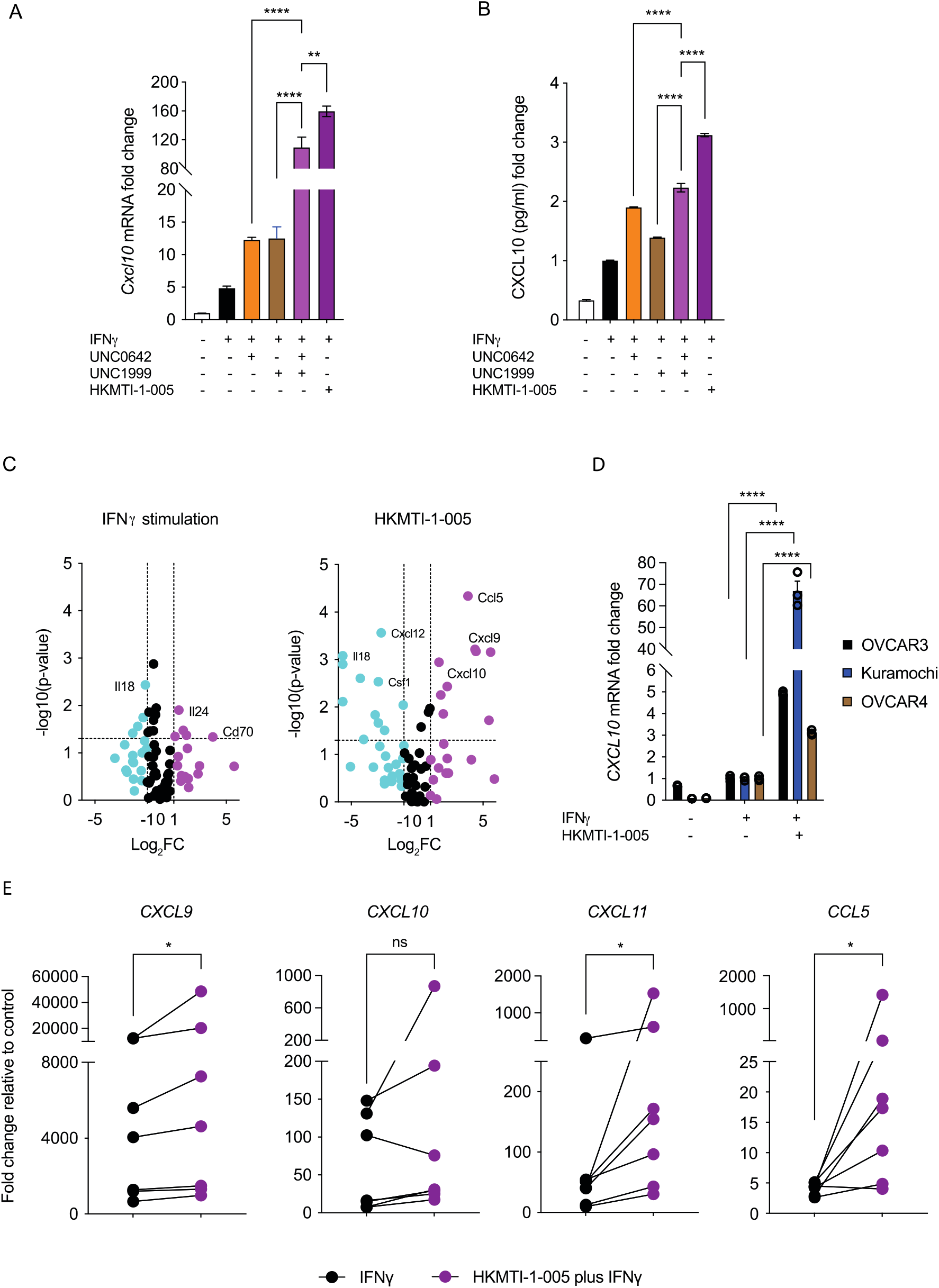
Dual inhibition of G9A/EZH2 upregulates chemotactic chemokines in vitro in mouse and human. A. *Cxcl10* mRNA fold change, normalised to *Gapdh* housekeeping gene, following treatment of *Trp53*^*-/-*^ID8 cells with IFN-γ 1ng/ml with or without UNC0642 5 μM (G9A inhibitor), UNC1999 2 μM (Ezh2 inhibitor) or HKMTI-1-005 6 μM (dual G9A/EZH2 inhibitor) for 72 hours. Mean values across treatments were compared with ordinary one-way ANOVA with Tukey’s multiple comparison test. Bars represents mean +/-SEM, n=3 biological replicates. B. CXCL10 protein fold change determined by ELISA following treatment as per A. Mean values across treatments were compared using one-way ANOVA with Dunnett’s multiple comparisons test. Bars represents mean +/-SEM, n=3 biological replicates. C. Expression of 84 chemokines and cytokines following treatment of *Trp53*^*-/-*^ID8 cells with IFN-γ 1ng/ml (left panel) with or without treatment with HKMTI-1-005 6 μM (right panel) for 72 hours. The experiment was performed in technical triplicates. Magenta colour: Log_2_FC ≥ 1, black colour: -1 ≤ Log_2_FC ≤ 1 and cyan colour: Log_2_FC ≤ -1 Gene list is found on Table S3, quality control and automatic normalization analysis in Table S4 and Table S5, respectively. D. *CXCL10* mRNA change following treatment of OVCAR3, Kuramochi and OVCAR4 cell lines with IFN-γ 1ng/ml or HKMTI-1-005 10 μM plus IFN-γ 1ng/ml for 72 hours. ΔΔC_T_ values were normalised to control *ACTB* housekeeping gene C_T_ values. Mean values across treatments were compared by ordinary one-way ANOVA with Tukey’s multiple comparison test and only comparisons between HKMTI-1-005 10 μM plus IFN-γ versus IFN-γ alone are shown. Each bar represents mean +/-SEM, n=3 biological replicates. E. *CXCL9, CXCL10, CXCL11* and *CCL5* expression in human ascites derived cultures. Each dot represents the median of 3 technical replicates for n=7 patients. ΔΔC_T_ values were normalised to *ACTB* housekeeping gene. C_T_ values. Median values were compared using the Wilcoxon matched-pair test. ****p <0.0001, ***p <0.001, **p <0.01, *p <0.05, ns= non-significant.

An 84 chemokine/cytokine gene expression array confirmed that dual G9A/EZH2 inhibition with HKMTI-1-005 had a potent effect on chemokine expression *in vitro*. Specifically, it upregulated *Cxcl10* 3-fold (p=0.001), *Cxcl9* 22-fold (p=0.0006) and *Ccl5* 14-fold (p<0.0001), compared to IFN-γ stimulation alone (**Fig 2C**). By contrast, chemokines secreted upon cell death, such as interleukin-1 (*Il-1*) or interleukin-18 (*Il-18*) (46, 47), were not increased (*Il-1a* fold-change 0.43, p=0.34, *Il-1b* fold-change 0.61, p=0.30 and *Il-18* fold-change 0.02, p<0.0001). This implies that the upregulation of the CXCR3-binding chemokines CXCL9, CXCL10 and CCL5 is not a cell death-associated stress response. HKMTI-1-005 treatment also upregulated *CXCL10* transcription in established human HGSC cell lines, including OVCAR3 (fold change 4.9, p<0.0001), OVCAR4 (fold change 3.1, p<0.0001) and Kuramochi (fold change 66.9, p<0.0001), when compared to IFN-γ stimulation alone (**Fig 2D**). Importantly, treatment of patient ascites-derived primary HGSC cells (including one patient with high-grade endometrioid ovarian carcinoma) with HKMTI-1-005 significantly increased *CXCL9* (p=0.01), *CXCL11* (p=0.01) and *CCL5* (p=0.03) mRNA levels, compared to IFN-γ stimulation alone (**Fig 2E**), demonstrating that dual G9A/EZH2 inhibition may have an immunostimulatory effect in the tumour microenvironment of patients with HGSC.

### Dual G9A/EZH2 inhibition alters transcription and chromatin conformation in vivo

We hypothesised that altered chromatin accessibility induced by G9A/EZH2 inhibition could explain the changes in gene expression. To investigate this, we used the Assay of Transposase-Accessible Chromatin with sequencing (ATAC-seq) and RNA sequencing (RNA-seq) on tumours harvested after treatment with HKMTI-1-005 *in vivo*. As described in the Methods, 6-week-old C57BL/6J female mice were injected IP with 5×10^6^ *Trp53*^*-/-*^ID8 cells/mouse and treated with 20 mg/kg HKMTI-1-005 IP twice/daily for 2 weeks, starting on day 21 post cell inoculation. At the end of the treatment, tumours were harvested and analysed by ATAC-seq and RNA-seq. Analysis of the ATAC-seq showed that there were more peaks representing areas of open chromatin in the HKMTI-1-005-treated samples compared to controls (**Fig 3A**); most of the peaks were in intergenic regions (58.4%). Approximately 33% of peaks were intronic and 6.5% were in promoter regions (**Fig 3B**). Peaks were present in genes involved in the activation pathways for *Cxcl9, Cxcl10 and Ccl5*, including *Stat1, Irf1*, nuclear factor NF-kappa-B p105 subunit (*Nfkb1*) and inhibitor of nuclear factor kappa-B kinase subunit beta (*Ikbkb*) (data not shown).

**Figure 3:**
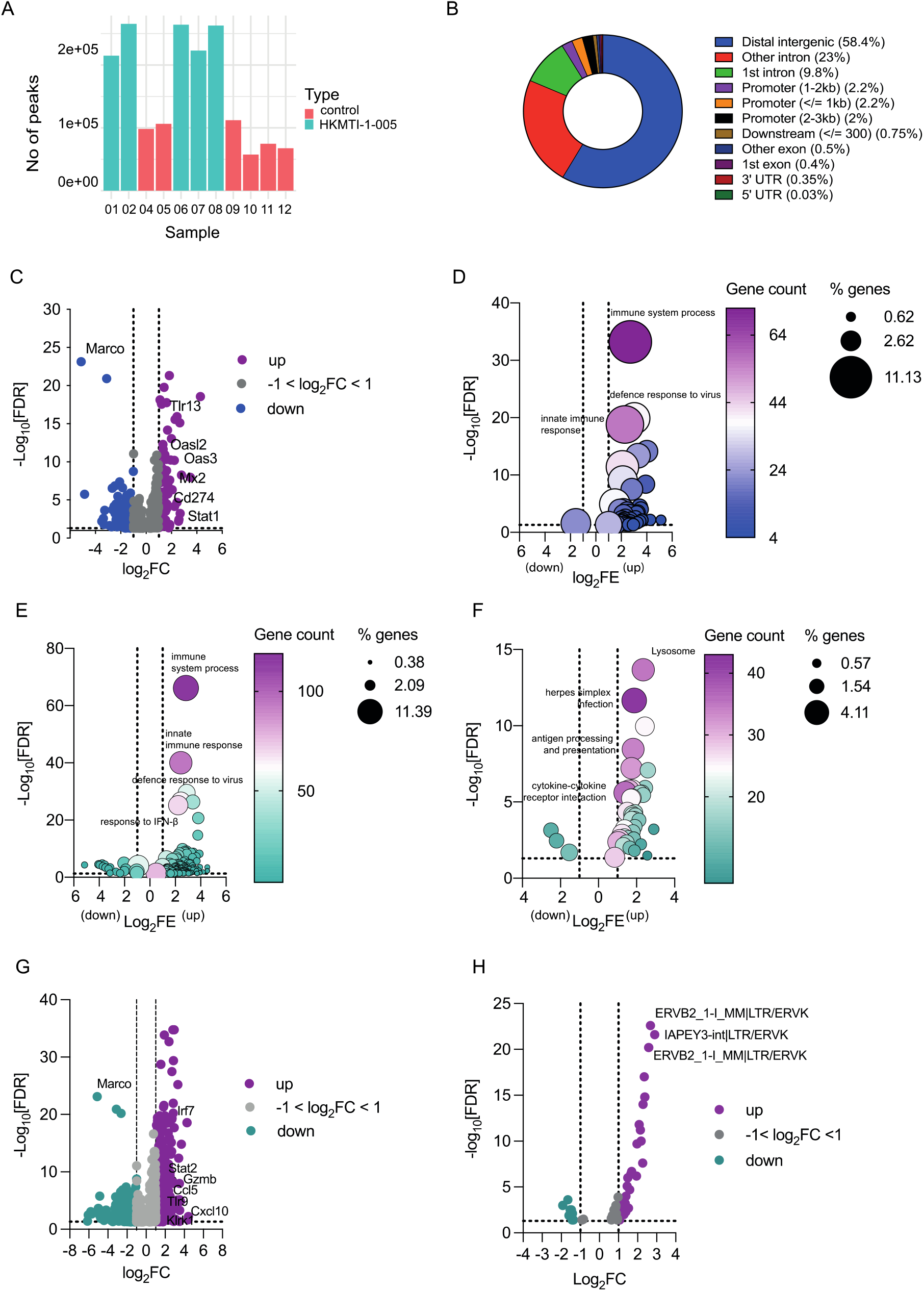
ATAC sequencing and RNA sequencing on omental tumour deposits. A. Number of ATACseq peaks for control (n=6) and HKMTI-1-005 samples (n=5), before applying filtration criteria. One HKMTI-1-005 sample did not yield enough sequencing reads and was removed from analysis. B. Distribution of ATAC seq peaks across the genome. C. Overlap of genes with ATACseq peaks showing increased chromatin accessibility that were also differentially expressed (n=1,106) by RNAseq following HKMTI-1-005 treatment *in vivo* (HKMTI-1-005 n=7, control n=7). FC = fold change. Purple colour: Log_2_FC ≥ 1, grey colour: -1 < Log_2_FC ≤ 1 and blue colour: Log_2_FC ≤ -1. D. Biological Processes (BP) sub-ontology for 1,053 genes from C that overlapped with gene expression signatures from Database for Annotation, Visualization and Integrated Discovery (DAVID) online Functional Annotation Tool. Gene count denotes the number of genes found to overlap with genes within the respective signature and the dot size represents the percentage of these genes within the signature. FE = fold enrichment, FDR = False discovery rate. E and F. Differentially expressed genes (DEG) following HKTMI-1-005 classified by BP and KEGG sub-ontologies, respectively. Gene count denotes the number of genes found to overlap with genes within the respective signature and the dot size represents the percentage of these genes within the signature. FE = fold enrichment, FDR = False discovery rate. G. Volcano plot showing individual DEG following HKMTI treatment (n=7) versus control (n=7) by RNA sequencing with an FDR <0.05. FC = fold change. Purple colour: Log_2_FC ≥ 1, grey colour: -1 < Log_2_FC ≤ 1 and green colour: Log_2_FC ≤ -1. H. Volcano plot showing differentially expressed Endogenous Retroviruses (ERVs), following HKMTI treatment (n=7) versus control (n=7). FC = fold change. Purple colour: Log_2_FC ≥ 1, grey colour: -1 < Log_2_FC ≤ 1 and green colour: Log_2_FC ≤ -1.

We found a statistically significant overlap of 1,106 genes in common between differentially expressed genes (DEG) identified by RNAseq and those in an euchromatin state identified by ATACseq (**Fig 3C**). Among these were the Toll-like receptor *Tlr13* (Log_2_FC 2.9, FDR= 7.42e-19), the IFN pathway mediator *Stat1* (Log_2_FC 1.01, FDR=4.1e-03), *Cd274*, which encodes PD-L1 (Log_2_FC 1.39, FDR=8.99e-06), and genes involved in antiviral response, such as *Mx2* (Log_2_FC 1.86, 2.43e-08) and *Oas3* (Log_2_FC 2.2, FDR=6.3e-11) (**Fig 3C**).

Using the DAVID functional annotation tool, we found that the most statistically significant upregulated genes with open chromatin belonged to signatures categorised as *immune system process* (GO:0002376, FDR=6.25e-34), *defence response to virus* (GO:0051607, FDR= 1.31e-20) and *innate immune response* (GO:0045087, FDR=1.58e-19), (**Fig 3D**). Other significantly upregulated pathways were *cellular response to IFN-β* (GO:0035458, FDR= 7.65e-15) and *response to virus* (GO:0009615, FDR=4.32e-14) (**Fig 3D**).

Treatment with HKMTI-1-005 also significantly altered the transcriptome of tumours *in vivo*. Functional annotation by biological processes sub-ontology showed that the most significantly upregulated pathway in the HKMTI-1-005-treated tumours was the immune pathway, *immune system process* (GO:0002376, FE 7.1, FDR=7.43e-67), with more than 10% DEGs overlapping with genes in this pathway (**Fig 3E**). *Innate immune response* (GO:0045087, FE 5.37, FDR=9.55e-41) and *defence response to virus* (GO:0051607, FE 7.4, FDR=3.12e-30) were also significantly upregulated (**Fig 3E**). Fewer pathways were downregulated following HKTMI-1-005. Amongst those were *positive regulation of glucose metabolic process* (GO:0010907, FE 13.6, FDR=1.82e-05), *positive regulation of lipid metabolic process* (GO:0045834, FE 16.6, FDR=3.21e-05) and *negative regulation of gluconeogenesis* (GO:0045721, FE 11.9, FDR=5.02e-05). Analysis using the KEGG database also revealed immune pathways being significantly enriched after treatment, such as *antigen processing and presentation* (mmu04612, FE 5.4, FDR=1.11e-10), *natural killer cell mediated cytotoxicity* (mmu04650, FE 3.3, FDR=6.45e-05) and *cytokine-cytokine receptor interaction* (mmu04660, FE 2.6, FDR=2.51e-06) (**Fig 3F**).

At the individual gene level, a total of 1,146 genes was upregulated and 733 genes downregulated following HKMTI-1-005 treatment (**Fig 3G**). Among the upregulated genes were *Cxcl10* (Log_2_FC 1.69, FDR<0.001), *Cxcl11* (Log_2_FC 1.19, FDR<0.001) and *Ccl5* (Log_2_FC 1.84, FDR= 8.33e-08). A number of other immune-stimulatory genes were also upregulated, including the gene encoding granzyme B (*GzmB)* (Log_2_FC 2.74, FDR=3.49e-09) and *Klrk1* (Log_2_FC 1.15, FDR=0.016), which encodes NKG2D, the major NK and T cell receptor for recognition and elimination of tumour cells (48). Moreover, *Stat2* (Log_2_FC 1.36, FDR= 1.61e-10) and *Tlr9* (Log_2_FC 1.21, FDR=5.00e-05), integral parts of type I IFN-mediated responses, were also upregulated. Other type I IFN system mediators were upregulated with treatment, such as *Irf7* (Log_2_FC 2.28, FDR=1.26e-19), *Irf9* (Log_2_FC 0.68, FDR=1.79e-05) and *Irf5* (Log_2_FC 0.41, FDR=0.02) (49), as well as type I IFN inducible genes including *Oasl1* (Log_2_FC 1.87, 3.61e-19), *Oas2* (Log_2_FC 1.94, 5.79e-11) and *Oas3* (Log_2_FC 2.22, 6.30e-11), which are all involved in the antiviral defence gene network (50). *Olfr732* showed the largest reduction (Log_2_FC -6.13, FDR=0.038), although its relation to cancer is unclear. Interestingly, the most statistically significant reduction occurred in Macrophage Receptor with Collagenous Structure *Marco*, (Log_2_FC -5.12, FDR=7.57e-24), the gene encoding a <u>pattern-recognition receptor</u> of the class A <u>scavenger receptor</u> family, expressed in tumour-associated macrophages (**Fig 3G**). ATACseq confirmed that *Marco* had reduced chromatic accessibility (**Fig 3C**), indicating HKMTI-1-005 may also act on tumour-associated macrophages.

Endogenous retroviruses (ERVs), ancient transposable elements integrated in the genome after infection, are epigenetically silenced under homeostatic conditions (51). Recent evidence suggests that they can potentiate anti-tumour immunity if transcriptionally active (6, 52, 53) and that, specifically in OC, ERV signatures can predict tumour lymphocyte infiltration and patient survival (54). Based on RNA-seq results indicating an upregulation of the *defence response to virus* pathway, we analysed the differential expression of ERVs following HKMTI-1-005 treatment. Out of 2,118 common ERVs detected between the HKMTI-1-005 treated and vehicle treated samples, 51 were differentially expressed at the 5% FDR threshold with 39/51 showing upregulation (**Fig 3H**). The IAP ERVK elements, IAPEY3-int|LTR/ERVK, had log_2_FC of 2.89 (FDR= 2.40e-22), whilst the ERVB2 ERVK elements had log_2_FC of 2.67 (FDR=2.41e-23). On the other hand, almost all of the downregulated retrotransposons in HKMTI-1-005-treated tumours belonged to the ERV1 class. The elements with the largest reduction were MuRRS4-int|LTR/ERV1 (log_2_FC -1.92, FDR<0.001) and MURVY-int|LTR/ERV1 (log_2_FC -1.6, FDR<0.001) (**Fig 3H**).

### Dual G9A/EZH2 inhibition prolongs survival in vivo

We next wanted to understand if modulating chromatin accessibility and stimulating gene expression, most importantly of chemokines associated with T and NK cell infiltration, could alter disease progression and/or the immune response in HGSC. The effect of HKMTI-1-005 on tumour growth was tested in mice bearing *Trp53*^*-/-*^ID8 tumours (omental and porta hepatis deposits) both by measuring tumour weight and ascites volume immediately after completion of treatment (**Fig 4A**) and also by measuring overall survival (**Fig 4B**). HKMTI-1-005 treatment resulted in a prolongation of median survival (48 *vs* 54.5 days, p <0.0001, HR 0.33, 95%CI 0.17-0.64) (**Fig 4C**). Treatment also resulted in a reduction of tumour weight at the end of treatment (135mg ± 5.2 *vs* 108mg ± 5.6, p=0.001, **Fig 4D**) and completely abrogated the development of ascites in this model (602 μl ± 297 μl *vs* 0 μl, p=0.0012, **Fig 4E**). There were no toxicity signals, over vehicle treatment, observed throughout treatment and no significant weight difference between treatment groups (**Fig S4**).

**Figure 4:**
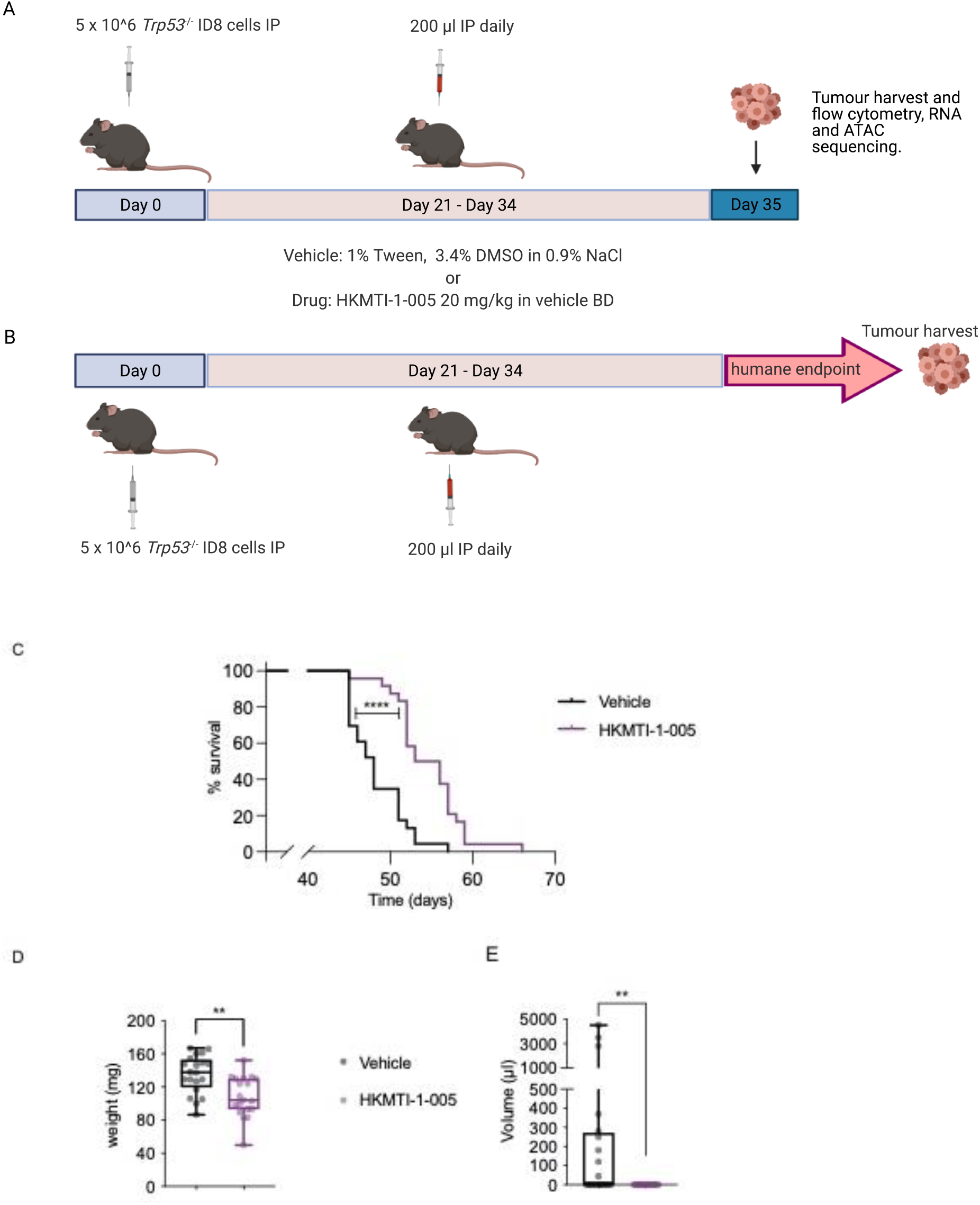
Dual G9A/EZH2 inhibition inhibits tumour growth and prolongs survival in a mouse ovarian cancer model. A. Experimental design for mechanism experiments. Mice bearing intraperitoneal *Trp53*^-/-^ID8 cells were treated with either HKMTI-1-005 (20mg/kg IP bd) or vehicle (1% Tween/3.4% DMSO in 0.9% NaCl IP bd) for 14 days starting on day 21, followed by omental and porta hepatic deposits harvest/weighting, measuring ascites and immunophenotyping by flow cytometry immediately after the end of treatment; Image created with BioRender.com B. Experimental design for efficacy experiments. Mice were treated as per (A) but treatment was followed by observation until mice reached humane survival endpoint. C. Kaplan-Meier survival curves for mice treated with vehicle (n=24) or 20 mg/kg HKMTI-1-005 (n=24) as per schedule on figure 4B. Median survival was 48 days for vehicle *vs* 54.5 days for HKMTI-1-005, p< 0.0001). Curves were compared using the Log-rank (Mantel-Cox) test, **** = p <0.0001. Experiment was performed twice with n=12 per cohort for each experiment. D. Whole tumour weight (including both porta hepatis and omental tumour deposits) and E ascites volume for mice treated with either vehicle (n=20) or HKMTI-1-005 (n=20) as per schedule in Fig 4A; comparisons were made using unpaired *t*-test for whole tumour burden and Mann-Whitney test for ascites volume, **p <0.01. Experiment was performed twice with n=10 per cohort for each experiment.

### Dual G9A/EZH2 inhibition alters immune composition in vivo in tumour and peritoneal cavity

To address whether the transcriptomic changes induced by the dual G9A/EZH2 inhibitor HKMTI-1-005 could stimulate an immune response *in vivo*, we examined the effect of treatment on the tumour microenvironment *in vivo*. C57BL/6J mice bearing *Trp53*^*-/-*^ID8 tumours were treated with HKMTI-1-005 20 mg/kg daily for 2 weeks (**Fig 4A**). At day 35, tumour deposits from two separate sites, omentum and porta hepatis, were immunophenotyped by flow cytometry. We hypothesised that the induction of chemokines (**Fig 3**) would stimulate immune cell infiltration. Accordingly, treatment with HKMTI-1-005 significantly increased the number of natural killer (NK) cells in both tumour sites (porta hepatis 3.8×10^6^ ± 0.8 *vs* 8.2×10^6^ ±1.8 cells/g, p=0.01; omentum 2.2×10^6^ ± 0.23 *vs* 3.6×10^6^ ± 0.39 cells/g, p=0.0067, **Fig 5A**). Moreover, the effector CD44^+^CD62L^-^CD8^+^ cytotoxic T cell population was significantly increased in the porta hepatis deposit (37.1% ± 11.5% *vs* 66.7% ± 4.9% effector CD8^+^/total CD8^+^, p=0.03) with a similar, but statistically non-significant trend in the omental deposit (67.2% ± 6.1% *vs* 75.04% ± 2.1%, p=0.27). Similarly, the naïve CD44^-^CD62L^+^ CD8^+^ T cell population was decreased in the porta hepatis (14.07% ± 4.5% *vs* 3.2% ± 1.6% naive CD8^+^/total CD8^+^, p=0.03) (**Fig 5B/C/D**). Additionally, the percentage of granzyme-B^+^ CD8^+^ cells was significantly higher following treatment in both tumour sites (porta hepatis 22.1% ± 6.6% *vs* 65.7% ± 2.7% GZM-B^+^CD8^+^/total CD8^+^, p<0.0001; omentum 32.3 ± 5.4% *vs* 64.59 ± 3.8% GZM-B^+^CD8^+^/total CD8^+^, p=0.0002) (**Fig 5E, F**), further indicating activation of effector CD8^+^ cells. Interestingly, these changes were accompanied by a decrease in the immunosuppressive FoxP3^+^ regulatory CD4^+^ population, mainly in omentum (1.2×10^6^ ± 0.16 *vs* 0.5×10^6^ ± 0.10 cells/g, p=0.014), with a statistically non-significant decrease in the porta hepatis (2.9×10^6^ ± 0.92 *vs* 1.2×10^6^ ± 0.15 cells/g, p=0.114) (**Fig 5G**).

**Figure 5:**
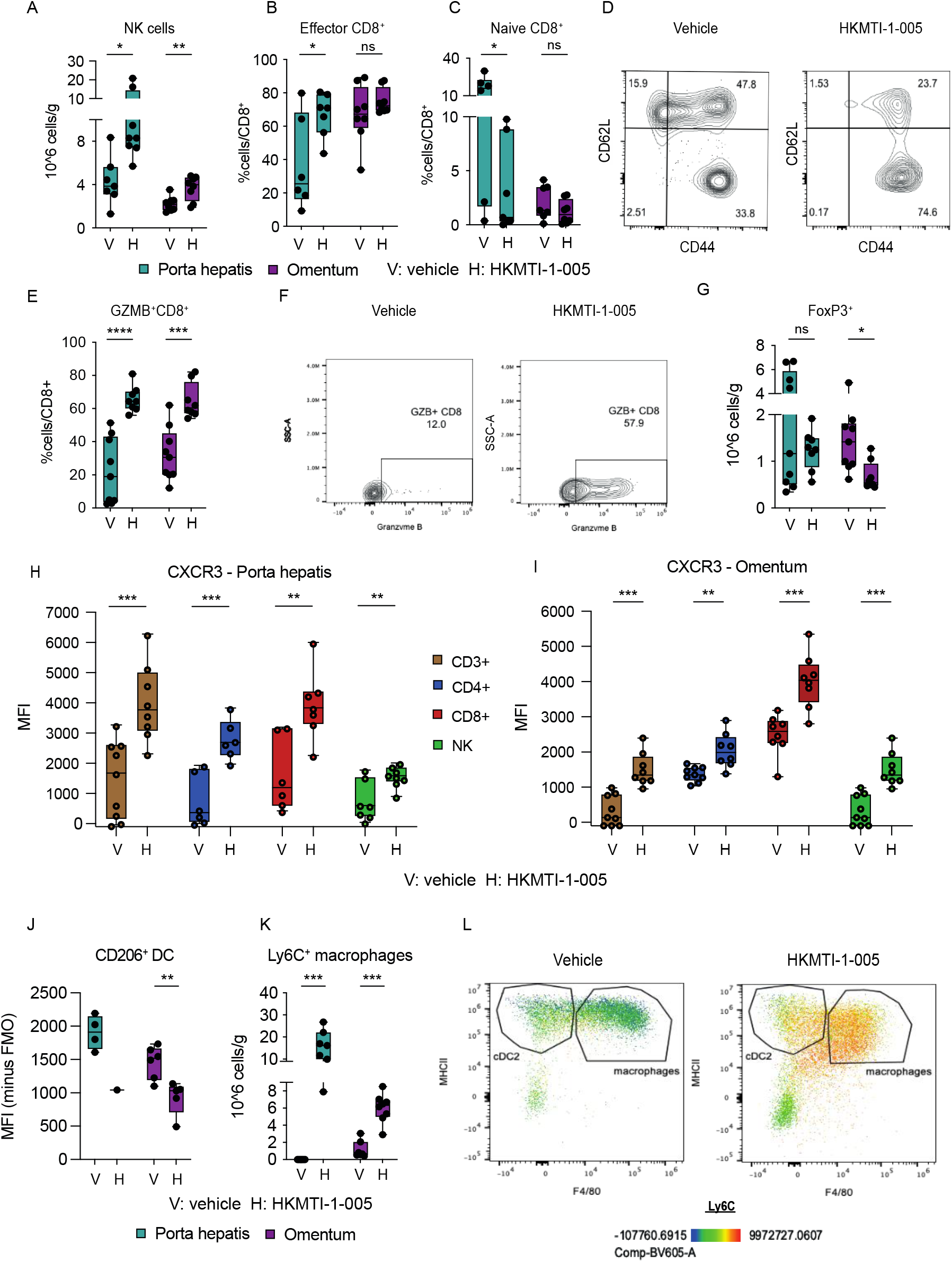
Dual G9A/EZH2 inhibition changes intra-tumoural immune cell composition n a mouse ovarian cancer model. A. Quantitative result for NK cells (CD3^-^ DX5^+^) cells in porta hepatis deposits with vehicle (n=7) *vs* HKMTI-1-005 (n=8) treatment and in the omental deposits in mice treated with vehicle (n=9) *vs* HKMTI-1-005 (n=8) treatment from mice bearing intraperitoneal *Trp53*^-^ID8 cells were treated with either HKMTI-1-005 or vehicle as per figure 4A. Significance was tested using an unpaired *t*-test. B. Percentage of effector CD8^+^ cells (CD44^+^CD62L^-^) within the total CD8^+^ population in porta hepatis deposits and omental deposits in mice treated with vehicle (n=6, 8 respectively) *vs* HKMTI-1-005 (n=7, 8 respectively) treatment. Significance was tested using an unpaired *t*-test test. C. Percentage of naïve CD8^+^ (CD44^-^ CD62L^+^) within the total CD8^+^ population in porta hepatis deposits and omental deposits in mice treated with vehicle (n=6, 8 respectively) *vs* HKMTI-1-005 (n=7, 8 respectively) treatment. Significance was tested using an unpaired *t*-test (porta hepatis) and Mann-Whitney test (omental deposit). D. Representative contour plot from one of the omental deposits for effector and naïve CD8^+^ cells, in mice treated with vehicle *vs* HKMTI-1-005 treatment. E. Percentage of granzyme-B (GZMB^+^) CD8^+^ cells, following stimulation, within the total CD8^+^ population in porta hepatis deposits and omental deposits from mice treated with vehicle (n=9, 8 respectively) *vs* HKMTI-1-005 (n=9, 8 respectively) treatment. Statistical significance was tested by unpaired *t*-test. F. Representative contour plot showing (GZMB^+^) CD8^+^ cells from one of the omental deposits from E. G. Quantitative result for T regulatory CD4^+^ cells (FoxP3^+^ CD4^+^) cells in porta hepatis deposits and omental deposits with vehicle (n=9, 8 respectively) *vs* HKMTI-1-005 (n=9, 8 respectively) treatment. Unpaired *t*-test (porta hepatis deposits) and Mann-Whitney test (omental deposits). H. CXCR3 median fluorescence intensity (MFI) on CD3^+^ (n=9 vehicle and n=8 HKMTI-1-005), CD4^+^ (n=6 vehicle and n=6 HKMTI-1-005), CD8^+^ (n=6 vehicle and n=7 HKMTI-1-005) and NK cells (n=7 vehicle and n=8 HKMTI-1-005) in the porta hepatis deposits. Statistical significance was tested by unpaired *t*-test. I. CXCR3 MFI on CD3^+^ (n=9 vehicle and n=8 HKMTI-1-005), CD4^+^ (n=9 vehicle and n=8 HKMTI-1-005), CD8^+^ (n=8 vehicle and n=8 HKMTI-1-005) and NK cells (n=9 vehicle and n=8 HKMTI-1-005) in omental deposits. Unpaired *t*-test. J. CD206 MFI on cDC1 dendritic cells (CD11b^-^ MHCII^+^CD11c^+^) in porta hepatis deposits (n=4 vehicle and n=1 HKMTI-1-005, statistics not performed as n<3) and omental deposits (n=6 vehicle and n=5 HKMTI-1-005). Statistical significance was tested by unpaired *t*-test. K. Ly6C^+^ macrophages (CD11b^+^MHCII^+^F4/80^+^) in porta hepatis and omentum deposits with vehicle (both n=7) *vs* HKMTI-1-005 (n=7, 8 respectively) treatment. Statistical significance was tested by the Mann-Whitney test. L. Representative flow cytometry plot with pseudocolour heatmap showing Ly6C^+^ macrophages from a representative omental deposit from K. cDC2+ cells were subsequently gated on a CD11c+. ****p <0.0001, ***p<0.001, **p <0.01, *p <0.05, ns= non-significant. Error bars represent SEM.

Expression of CXCR3 was increased on all lymphoid subsets in both tumour deposits, complementing our *in vitro* data (porta hepatis 1581 ± 504.3 *vs* 3899 ± 432.3 MFI on CD8^+^ cells, p=0.0049; omentum 2485 ± 204.0 *vs* 4012 ± 273.6 MFI on CD8^+^ cells, p=0.0005, **Fig 5H, I**).

We further observed that HKMTI-1-005 treatment decreased expression of CD206 receptor on cDC1 (omental tumour 1456 ± 100.5 *vs* 925.8 ± 113.6 MFI, p=0.006), a marker mainly associated with induction of T cell tolerance (55) (**Fig 5J**). CD206 was also under-expressed in macrophages in the omental deposit (5220 ± 508 *vs* 7620 ± 626 MFI, p=0.01, data not shown), further supporting the hypothesis that G9A/EZH2 inhibition can block steps that lead to immunosuppression (56) and hence, create a microenvironment that is conducive to cytotoxic T cell responses. HKMTI-1-005 treatment also resulted in a profound increase of Ly6C^+^ macrophages compared to vehicle (porta hepatis 0×10^6^ *vs* 16.21×10^6^ ± 2.56 cells/g, p=0.0006; omentum 0.64 ± 0.38 ×10^6^ *vs* 6.32×10^6^ ± 0.59 cells/g, p=0.0006, **Fig 5K, L**). Ly6C is a marker mostly expressed by precursors of tumour associated macrophages (TAMs) and is thought to be downregulated as these precursors differentiate into TAMs (57). Immunohistochemistry staining confirmed similar trends in the omentum with regards to NK and FoxP3^+^ cell populations (**Fig S5**).

The peritoneal cavity is an important site for the transcoelomic spread of HGSC with many patients developing ascites, which is rich in both tumour and immune cells. In the *Trp53*^*-/-*^ID8 model, G9A/EZH2 blockade increased the absolute number of peritoneal NK cells (12.8×10^3^ ± 2.45 *vs* 36.9×10^3^ ± 6.66 cells/ml, p=0.006) (**Fig 6A, C**) and a higher percentage of IFN-γ^+^ NK cells, indicative of an active anti-tumour response (27.5% ± 7.29 *vs* 62.5% ± 3.97 IFN-γ^+^NK cells/total NK cells, p=0.001) (**Fig 6B**). As with the solid tumour deposits, we observed a depletion of T regulatory cells in the peritoneal cavity (2.5×10^3^ ± 0.55 *vs* 0.7×10^3^ ± 0.29 cells/ml, p=0.01) (**Fig 6D/F**) and an increase in Ly6C^+^ macrophages (10.7×10^3^ ± 3.4 *vs* 51.6×10^3^ ± 7.62 cells/ml, p=0.0005) (**Fig 6E**). Furthermore, mirroring our intratumoral findings, the expression of CXCR3 was significantly increased on all lymphoid cell subpopulations (1746 ± 318 *vs* 6144 ± 460 MFI on CD8^+^ cells, p<0.0001) (**Fig 6G, H**) suggesting that this response is driven by the CXCL10 axis.

**Figure 6:**
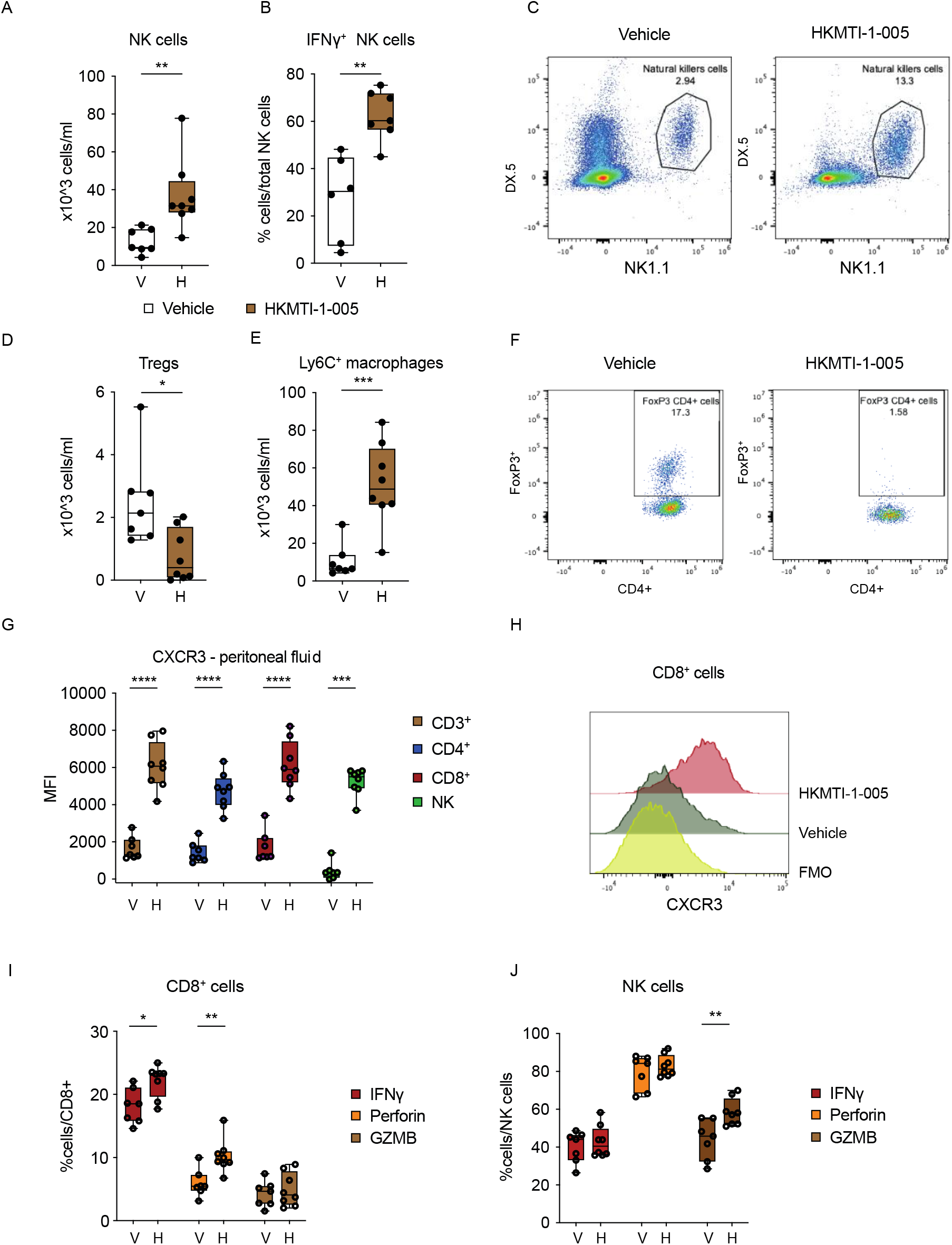
Dual G9A/EZH2 inhibition changes peritoneal cavity immune cell composition and chemokine milieu in the spleen in a mouse ovarian cancer model. A. Density of NK cells (CD3^-^ NK1.1^+^DX5^+^) in the peritoneal fluid with vehicle (n=7) *vs* HKMTI-1-005 (n=8) treatment from mice bearing intraperitoneal *Trp53*^-/-^ID8 cells as per figure 4A. Statistical significance was tested by unpaired *t*-test. B. Percentage IFN-γ^+^ NK cells in peritoneal fluid from mice treated with vehicle (n=6) *vs* HKMTI-1-005 (n=7) treatment. Statistical significance was tested by unpaired *t*-test. C. Representative flow cytometry plot from A. D. Density of regulatory T cells (CD4^+^FoxP3^+^) in peritoneal fluid from mice treated with vehicle (n=7) *vs* HKMTI-1-005 (n=8) treatment. Statistical significance was tested by unpaired *t*-test. E. Density of Ly6C^+^ cells in the peritoneal fluid from mice treated with vehicle (n=7) *vs* HKMTI-1-005 (n=8) treatment. Statistical significance was tested by unpaired *t*-test. F. Representative flow cytometry plot from D. G. CXCR3 MFI on CD3^+^ (n=7 vehicle and n=8 HKMTI-1-005), CD4^+^ (n=7 vehicle and n=8 HKMTI-1-005), CD8^+^ (n=7 vehicle and n=8 HKMTI-1-005) and NK cells (n=7 vehicle and n=8 HKMTI-1-005) in the peritoneal fluid. Unpaired *t*-test was used to compare all mean values except NK cell comparison where the Mann-Whitney test was used. H. Representative MFI histogram on one sample from G; FMO= fluorescence-minus-one. I and J. Percentage of IFNγ, perforin and GZMB+ CD8^+^ cells (I) and NK cells (J) within their respective overall populations in spleen, following *ex vivo* stimulation with PMA and ionomycin and protein transport inhibitors brefeldin and monensin in mice treated with either vehicle (n=7) *vs* HKMTI-1-005 (n=8) treatment. Statistical significance was tested by unpaired *t*-test. ****p <0.0001, ***p<0.001, **p <0.01, *p <0.05, ns= non-significant. Error bars represent SEM.

Finally, we analysed the spleens of mice following G9A/EZH2 blockade to determine if treatment could potentiate an adaptive T cell response in this important secondary lymphoid organ. Interestingly, we found that spleens treated with HKMTI-1-005 contained a higher percentage of CD8^+^ cells containing intracellular IFN-γ (18.1% ± 1.04% *vs* 22.1% ± 0.9% IFN-γ^+^CD8^+^/total CD8^+^ cells, p=0.012) and perforin (5.9% ± 0.8% *vs* 10.2% ± 0.9% perforin positive CD8^+^/total CD8^+^ cells, p=0.004, **Fig 6I**). We also observed a higher percentage of NK cells containing granzyme-B (43.9% ± 4.0% *vs* 58.4% ± 2.5% GZMB positive NK/total NK cells, p=0.007, **Fig 6J**). This provided further evidence that G9A/EZH2 blockade drives an anti-tumoural response, not only directly in the tumour deposits but also more systemically.

## Discussion

In this study, we investigated whether modulation of epigenetic pathways could potentiate the immune response against HGSC, using a mouse model that recapitulates many aspects of the human disease. Dual blockade of the histone methyltransferases G9A and EZH2 reprogrammed the immune tumour microenvironment (TME) and activated the transcription of immune networks *in vivo*. This led to alteration of the immune cell composition in the TME, with accumulation of effector cytotoxic lymphocytes and NK cells, whilst reducing the population of immunosuppressive Treg CD4^+^ cells. We observed that G9A/EZH2 inhibition activated the CXCR3 axis, with tumour cell-derived IFN-γ-inducible chemokines CXCL9, CXCL10, CXCL11 and CCL5 being upregulated *in vitro* and *in vivo*, with a corresponding increase in the expression of receptor CXCR3 on immune cells *in vivo*. Importantly, HKMTI-1-005 treatment inhibited tumour growth and prolonged survival *in vivo*. The chemokine induction by HKMTI-1-005 was also observed in established human cell lines and ascites-derived primary spheroids, making this an attractive inhibitor for potential future clinical application.

We found that treatment increased cytotoxic CD8^+^ T cell number in the spleen following treatment. Furthermore, treatment also reduced the expression of the suppressive receptors CD206 on dendritic cells and macrophages, and blocked monocyte-to-macrophage differentiation in both the TME and peritoneal cavity. It is widely recognised that TAMs derive from the large population of CCR2^high^Ly6C^+^ inflammatory monocytes that constantly contributes to the pool, and that Ly6C expression gradually reduces as TAMs differentiate within tumours (58, 59). HKMT-1-005 treatment increased the abundance of Ly6C^+^ macrophages, suggesting that this epigenetic modifier may impede the differentiation of the monocyte precursor pool into fully differentiated TAMs.

The preclinical results presented here provide evidence that dual inhibition of G9A and EZH2 induces more robust chemokine induction than blockade of either methyltransferase alone. Recently, the co-dependence of EZH2 and G9A was established by Mozzetta et al (45) and this has led to efforts of discovering pharmacological inhibitors that target both enzymes simultaneously, with HKMTI-1-005 being the first one described (30). Curry et al. showed that treatment of the breast cancer cell line MDA-MB-231 with HKMTI-1-005 induced transcription of *SPINK1*, which does not occur EZH2 or G9A were knocked down (30), strongly suggesting that the two enzymes are co-dependent in silencing certain genes. Novel EZH2 inhibitors have already entered early phase clinical trials in solid tumours, and tazemetostat was granted FDA approval for use in EZH2-mutated refractory/relapsed follicular lymphoma in 2020 (60).

This work has generated interesting questions with regards to mechanism of action of G9A/EZH2 blockade that will need further investigation. Firstly, our transcriptional and chromatin accessibility analyses were based on whole-tumour sequencing and therefore do not specify the cell type subjected to transcriptional modifications by HKMTI-1-005 treatment. Single-cell sequencing may help to delineate the exact cell type(s) on which the compound exerts its function in the TME. Secondly, the contribution of ERV-K retroelements in the initiation of immune responses following HKMTI-1-005 treatment warrants further exploration. Recent evidence suggests that ERVs can potentiate anti-tumour immunity when they are transcriptionally active (6, 52, 53) and that the activation of evolutionary young elements is associated with innate immune responses (61). ERVK, an evolutionary young element, was activated following treatment with HKMTI-1-005 and, interestingly, antibodies against ERVK have been detected in the serum of OC patients (62). The downregulation of the macrophage receptor MARCO following HKMTI-1-005 treatment is also an intriguing finding; inhibiting MARCO reprogrammes macrophages to acquire an anti-tumour phenotype, inhibiting tumour cell growth (63, 64). In the work presented here, we used a single ovarian cancer mouse model, the ID8 mouse model, engineered with *Trp53*^-/-^deletion, which represents the most common genetic aberration in human ovarian HGSC. This model reflects the intrabdominal dissemination of OC with peritoneal deposits and ascites, as commonly observed in human disease, and it has been used to study alterations of the immune cell composition in the tumour microenvironment (65, 66). To reinforce our findings with the *Trp53*^-/-^ID8 model, we used established human cell lines and primary HGSC spheroids.

Overall, the results of this work support the hypothesis that dual blockade of G9A/EZH2 histone methyltransferases modulates anti-tumour immune responses, confers a survival benefit in an aggressive murine model of OC and warrants further investigation towards clinical development.

## Funding and acknowledgments

This work was funded by the NIHR Imperial Biomedical Research Centre (grant reference P74580, P74580-2 and P77646 – these grants did not support any animal experiments), Ovarian Cancer Action (grant reference 006), Cancer Research UK (grant references A17196, A29799, A15973, A17196, A12295) and the University of Glasgow endowments. MJF would like to thank the Engineering and Physical Science Research Council for an Established Career Fellowship (EP/R00188X/1)

Infrastructure support was provided by the Cancer Research UK/NIHR Imperial Experimental Cancer Medicine Centre, the Imperial College Healthcare Tissue Bank and the CRUK Imperial and Glasgow Centres.

The authors would like to thank the LMS/NIHR Imperial Biomedical Research Centre Flow Cytometry Facility and the Biological Services and Histology Service at the CRUK Beatson Institute for their support.

The authors would also like to acknowledge the support of Dr Seth Coffelt, Dr Josephine Walton and Dr Catherine Winchester.

