## Supplementary Figures and Methods for "Dual G9A and EZH2 inhibition stimulates an anti-tumour immune response in ovarian high-grade serous carcinoma": Supplementary Figures and Methods.docx

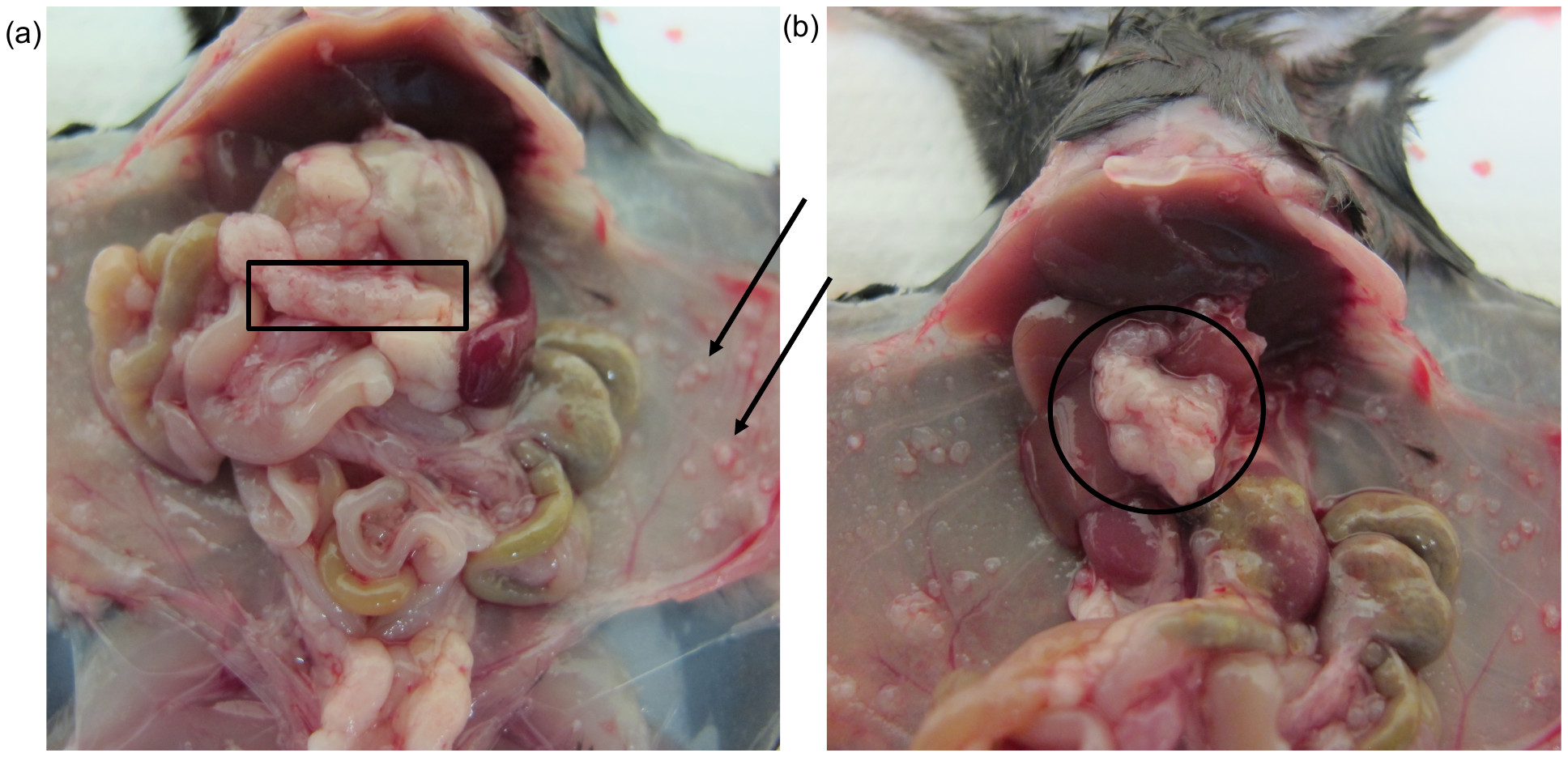

**Figure S1: Intra-abdominal tumour deposits in C57BL/6J mice following intraperitoneal inoculation with the *Trp53^-/-^* ID8 mouse model**

(a) C57BL/6J mouse; omental deposit (rectangle) and peritoneal deposits (arrows) are sites of disease, 6 weeks after IP inoculation with 5x10^6^ *Trp53^-/-^* ID8 cells in 200 μl of PBS. (b) Porta hepatis deposit (circle) after part of liver removed.

| Gene symbol | Exon spanning region | Catalogue Number |
| --- | --- | --- |
| Actb (mouse) | 2-3 | Mm02619580_g1 |
| ACTB (human) | 1-3 | Hs01060665_g1 |
| Cxcl10 (mouse) | 1-2 | [Mm00445235_m1](https://www.thermofisher.com/taqman-gene-expression/product/Mm00445235_m1?CID=&ICID=&subtype=) |
| CXCL10 (human) | 1-2 | Hs00171042_m1 |
| 18S ribosomal RNS (Rn18s, mouse) | - | [Mm03928990_g1](https://www.thermofisher.com/taqman-gene-expression/product/Mm03928990_g1?CID=&ICID=&subtype=) |
| GAPDH (human) | 6-8 | Hs02786624_g1 |
| RANTES (CCL5, human) | 1-2 | Hs00982282_m1 |
| Rantes (Ccl5, mouse) | 1-2 | Mm01302427_m1 |
| MIP3b (Ccl19, mouse) | 1-2 | Mm00839966_g1 |
| MCP-5 (Ccl12, mouse) | 1-2 | Mm01617100_m1 |
| Stat1 (mouse) | 20-21, 23-24 | Mm01257286_m1 |
| CXCL11 (human) | 1-2 | Hs00171138_m1 |
| MIP-3a (CCL20, human) | 2-3 | Hs00355476_m1 |

**Table S1: Primers used for single-gene RT-qPCR.**

All primers supplied by ThermoFisher.

| **Probe Name** | **Family** | **Target** |
| --- | --- | --- |
| **LAQ824** | HDAC - class I, IIa, IIb | HDAC 1/p21 promoter activation |
| **UNC0638** | Methyltransferase | G9a, GLP |
| **A-366** | Methyltransferase | G9a (EHMT2), GLP |
| **PFI-4** | Bromodomains | BRPF1B |
| **SGC0946** | Methyltransferase | DOTL-1 |
| **UNC0642** | Methyltransferase | G9a, GLP |
| **GSK343** | Methyltransferase | EZH2 |
| **GSK2801** | Bromodomains | BAZ2A/2B |
| **IOX2** | Oxyglutarate oxygenase | PHD2 |
| **NI-57** | Bromodomains | BRPF1, BRPF2, BRPF3 |
| **LLY507** | Methyltransferase | SMYD2 |
| **GSK484** | Arginine deiminase | PAD-4 |
| **PFI-3** | Bromodomains | SMARCA, PB1 |
| **UNC1215** | Methyl-lysine binder | L3MBTL3 |
| **I-CBP112** | Bromodomains | CREBBP, EP300 |
| **BAZ2-ICR** | Bromodomains | BAZ2A, BAZ2B |
| **SGC 707** | Methyltransferase | PRMT3 |
| **MS049** | Methyltransferase | PRMT4,6 |
| **NVS-1** | Bromodomains | CECR2 |
| **KDOAM25** | Demethylase | KDM5 |
| **GSK591** | Methyltransferase | PRMT5 |
| **BAY-598 (S-4)** | Methyltransferase | SMYD2 |
| **OICR9429** | Methyltransferase | WDR5 |
| **A196** | Methyltransferase | SUV420 H1/H2 |
| **MS023** | Methyltransferase | Type I PRMTs |
| **IOX1** | Oxyglutarate oxygenase | pan-2-OG |
| **OF-1** | Bromodomains | BRPF1, BRPF2, BRPF3 |
| **IBRD9** | Bromodomains | BRD9 |
| **LP99** | Bromodomains | BRD9/BRD7 |
| **SGC-CBP30** | Bromodomains | CREBBP, EP300 |
| **PFI-2** | Methyltransferase | SETD7 |
| **BI-9564** | Bromodomains | BRD9/BRD7 |
| **GSK-LSD1** | Demethylase | LSD-1 |
| **C646** | HAT | p300/CBP |
| **Bromosporine** | Bromodomains | Pan-Bromodomain |
| **PFI-1** | Bromodomains | BRD2, BRD3, BRD4, BRDT (BET) |
| **(+) JQ1** | Bromodomains | BET (BRD2-4 and BRDT) |
| **GSK-J4** | Histone Demethylase | JMJD3/UTX |
| **CI-994** | Histone deacetylase | HDAC - class I |
| **UNC1999** | Methyltransferase | EZH2 |

**Table S2: Library of epigenetic probes used in the drug-screening as per Structural Genomics Consortium (SGC, 2017).**

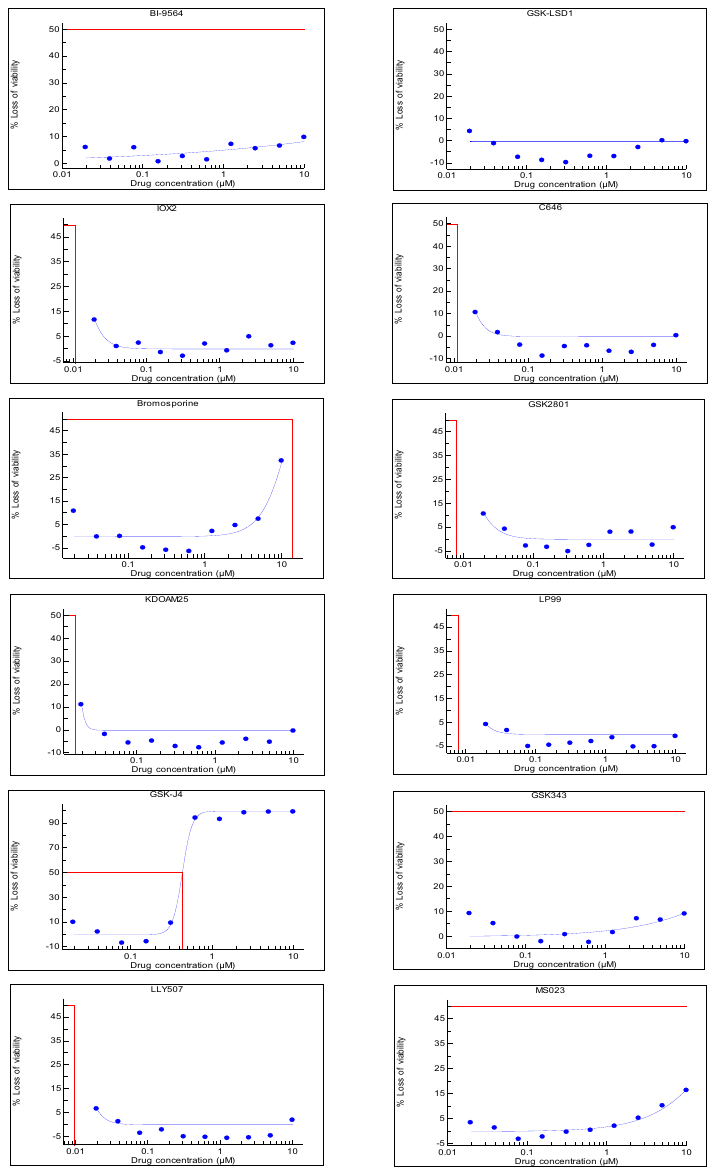

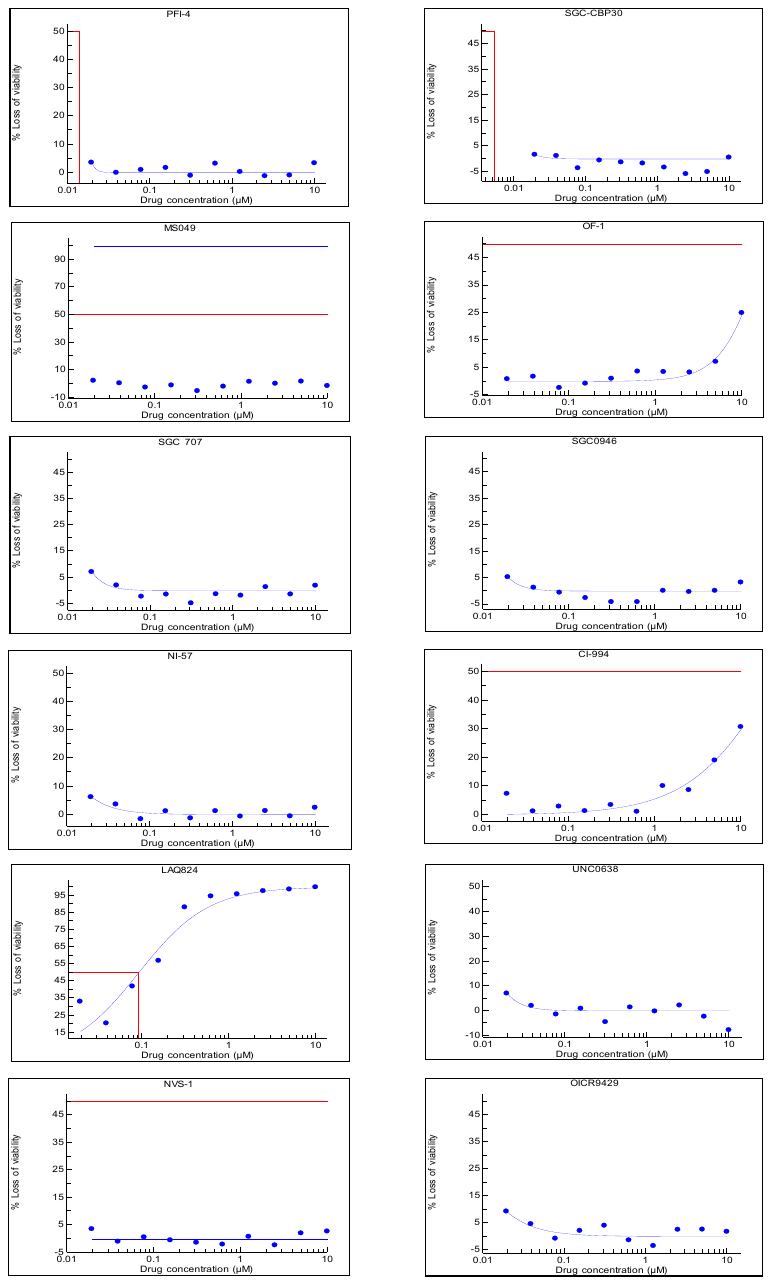

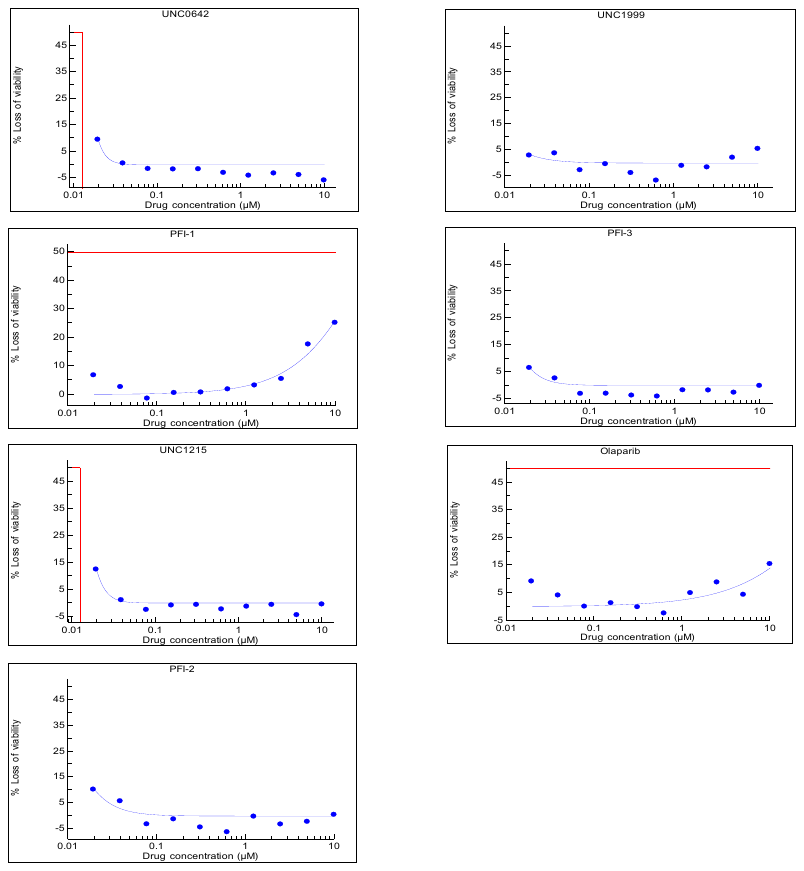

**Figure S2: Drug response curves for SGC library.**

2x10^3^ ID8 *Trp53*^-/-^ cells were plated on day 0 in 384 well plates and treated with the SGC library on day 1, at concentrations 20 nM - 10 μM. Loss of viability on y axis as measured by DAPI stained nuclei in treated wells relative to DMSO controls (n=3 technical replicates).

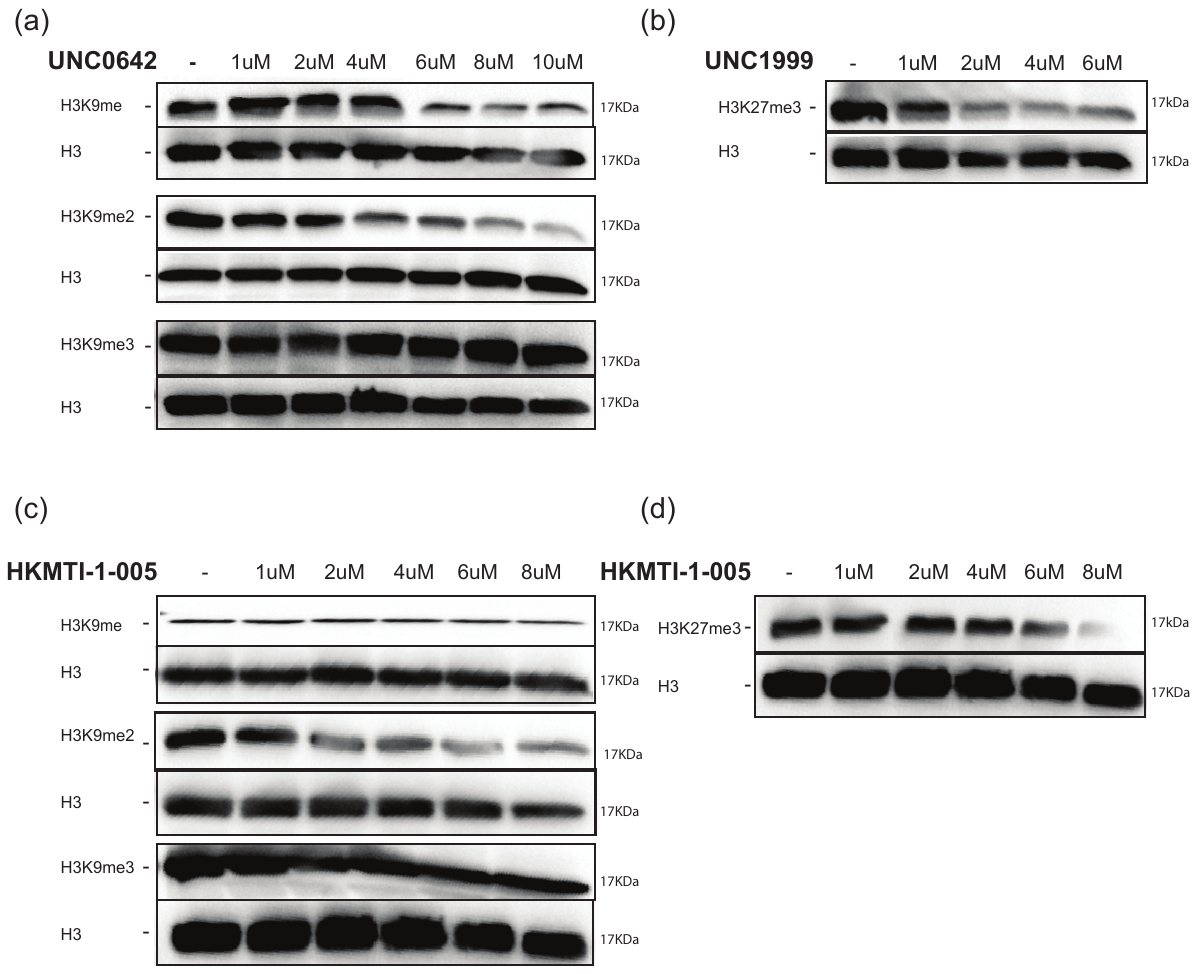
**Figure S3: Western blot analysis following inhibition of G9a, EZH2 and combination G9a/EZH2**.

Western blot analysis for H3K9 methylation marks with the G9a inhibitor, UNC0642 (a) and G9a/Ezh2 inhibitor, HKMTI-1-005 (c) and H3K27me trimethylation marks for Ezh2 inhibitor, UNC1999 (b) and HKMTI-1-005 (d). Histone H3 was used as a loading control. Protein electrophoresis was performed for each methylation mark and its respective loading control on separate gels, given that both the size of H3 and the size of the methylated mark were both approximately 17kDa.

| Position | RefSeq Number | Symbol | Description |
| --- | --- | --- | --- |
| A01 | NM_009605 | Adipoq | Adiponectin, C1Q and collagen domain containing |
| A02 | NM_007553 | Bmp2 | Bone morphogenetic protein 2 |
| A03 | NM_007554 | Bmp4 | Bone morphogenetic protein 4 |
| A04 | NM_007556 | Bmp6 | Bone morphogenetic protein 6 |
| A05 | NM_007557 | Bmp7 | Bone morphogenetic protein 7 |
| A06 | NM_011329 | Ccl1 | Chemokine (C-C motif) ligand 1 |
| A07 | NM_011330 | Ccl11 | Chemokine (C-C motif) ligand 11 |
| A08 | NM_011331 | Ccl12 | Chemokine (C-C motif) ligand 12 |
| A09 | NM_011332 | Ccl17 | Chemokine (C-C motif) ligand 17 |
| A10 | NM_011888 | Ccl19 | Chemokine (C-C motif) ligand 19 |
| A11 | NM_011333 | Ccl2 | Chemokine (C-C motif) ligand 2 |
| A12 | NM_016960 | Ccl20 | Chemokine (C-C motif) ligand 20 |
| B01 | NM_009137 | Ccl22 | Chemokine (C-C motif) ligand 22 |
| B02 | NM_019577 | Ccl24 | Chemokine (C-C motif) ligand 24 |
| B03 | NM_011337 | Ccl3 | Chemokine (C-C motif) ligand 3 |
| B04 | NM_013652 | Ccl4 | Chemokine (C-C motif) ligand 4 |
| B05 | NM_013653 | Ccl5 | Chemokine (C-C motif) ligand 5 |
| B06 | NM_013654 | Ccl7 | Chemokine (C-C motif) ligand 7 |
| B07 | NM_011616 | Cd40lg | CD40 ligand |
| B08 | NM_011617 | Cd70 | CD70 antigen |
| B09 | NM_170786 | Cntf | Ciliary neurotrophic factor |
| B10 | NM_007778 | Csf1 | Colony stimulating factor 1 (macrophage) |
| B11 | NM_009969 | Csf2 | Colony stimulating factor 2 (granulocyte-macrophage) |
| B12 | NM_009971 | Csf3 | Colony stimulating factor 3 (granulocyte) |
| C01 | NM_007795 | Ctf1 | Cardiotrophin 1 |
| C02 | NM_009142 | Cx3cl1 | Chemokine (C-X3-C motif) ligand 1 |
| C03 | NM_008176 | Cxcl1 | Chemokine (C-X-C motif) ligand 1 |
| C04 | NM_021274 | Cxcl10 | Chemokine (C-X-C motif) ligand 10 |
| C05 | NM_019494 | Cxcl11 | Chemokine (C-X-C motif) ligand 11 |
| C06 | NM_021704 | Cxcl12 | Chemokine (C-X-C motif) ligand 12 |
| C07 | NM_018866 | Cxcl13 | Chemokine (C-X-C motif) ligand 13 |
| C08 | NM_023158 | Cxcl16 | Chemokine (C-X-C motif) ligand 16 |
| C09 | NM_203320 | Cxcl3 | Chemokine (C-X-C motif) ligand 3 |
| C10 | NM_009141 | Cxcl5 | Chemokine (C-X-C motif) ligand 5 |
| C11 | NM_008599 | Cxcl9 | Chemokine (C-X-C motif) ligand 9 |
| C12 | NM_010177 | Fasl | Fas ligand (TNF superfamily, member 6) |
| D01 | NM_008155 | Gpi1 | Glucose phosphate isomerase 1 |
| D02 | NM_010406 | Hc | Haemolytic complement |
| D03 | NM_010503 | Ifna2 | Interferon alpha 2 |
| D04 | NM_008337 | Ifng | Interferon gamma |
| D05 | NM_010548 | Il10 | Interleukin 10 |
| D06 | NM_008350 | Il11 | Interleukin 11 |
| D07 | NM_008351 | Il12a | Interleukin 12A |
| D08 | NM_001303244 | Il12b | Interleukin 12b |
| D09 | NM_008355 | Il13 | Interleukin 13 |
| D10 | NM_008357 | Il15 | Interleukin 15 |
| Position | **RefSeq Number** | **Symbol** | **Description** |
| D11 | NM_010551 | Il16 | Interleukin 16 |
| D12 | NM_010552 | Il17a | Interleukin 17A |
| E01 | NM_145856 | Il17f | Interleukin 17F |
| E02 | NM_008360 | Il18 | Interleukin 18 |
| E03 | NM_010554 | Il1a | Interleukin 1 alpha |
| E04 | NM_008361 | Il1b | Interleukin 1 beta |
| E05 | NM_031167 | Il1rn | Interleukin 1 receptor antagonist |
| E06 | NM_008366 | Il2 | Interleukin 2 |
| E07 | NM_021782 | Il21 | Interleukin 21 |
| E08 | NM_016971 | Il22 | Interleukin 22 |
| E09 | NM_031252 | Il23a | Interleukin 23, alpha subunit p19 |
| E10 | NM_053095 | Il24 | Interleukin 24 |
| E11 | NM_145636 | Il27 | Interleukin 27 |
| E12 | NM_010556 | Il3 | Interleukin 3 |
| F01 | NM_021283 | Il4 | Interleukin 4 |
| F02 | NM_010558 | Il5 | Interleukin 5 |
| F03 | NM_001314054 | Il6 | Interleukin 6 |
| F04 | NM_008371 | Il7 | Interleukin 7 |
| F05 | NM_008373 | Il9 | Interleukin 9 |
| F06 | NM_008501 | Lif | Leukemia inhibitory factor |
| F07 | NM_010735 | Lta | Lymphotoxin A |
| F08 | NM_008518 | Ltb | Lymphotoxin B |
| F09 | NM_010798 | Mif | Macrophage migration inhibitory factor |
| F10 | NM_010834 | Mstn | Myostatin |
| F11 | NM_013611 | Nodal | Nodal |
| F12 | NM_001013365 | Osm | Oncostatin M |
| G01 | NM_019932 | Pf4 | Platelet factor 4 |
| G02 | NM_023785 | Ppbp | Pro-platelet basic protein |
| G03 | NM_009263 | Spp1 | Secreted phosphoprotein 1 |
| G04 | NM_009367 | Tgfb2 | Transforming growth factor, beta 2 |
| G05 | NM_009379 | Thpo | Thrombopoietin |
| G06 | NM_013693 | Tnf | Tumour necrosis factor |
| G07 | NM_008764 | Tnfrsf11b | Tumour necrosis factor receptor superfamily, member 11b (osteoprotegerin) |
| G08 | NM_009425 | Tnfsf10 | Tumour necrosis factor (ligand) superfamily, member 10 |
| G09 | NM_011613 | Tnfsf11 | Tumour necrosis factor (ligand) superfamily, member 11 |
| G10 | NM_033622 | Tnfsf13b | Tumour necrosis factor (ligand) superfamily, member 13b |
| G11 | NM_009505 | Vegfa | Vascular endothelial growth factor A |
| G12 | NM_008510 | Xcl1 | Chemokine (C motif) ligand 1 |
| H01 | NM_007393 | Actb | Actin, beta |
| H02 | NM_009735 | B2m | Beta-2 microglobulin |
| H03 | NM_008084 | Gapdh | Glyceraldehyde-3-phosphate dehydrogenase |
| H04 | NM_010368 | Gusb | Glucuronidase, beta |
| H05 | NM_008302 | Hsp90ab1 | Heat shock protein 90 alpha (cytosolic), class B member 1 |
| H06 | SA_00106 | MGDC | Mouse Genomic DNA Contamination |
| H07 | SA_00104 | RTC | Reverse Transcription Control |
| H08 | SA_00104 | RTC | Reverse Transcription Control |
| H09 | SA_00104 | RTC | Reverse Transcription Control |
| Position | **RefSeq Number** | **Symbol** | **Description** |
| H10 | SA_00103 | PPC | Positive PCR Control |
| H11 | SA_00103 | PPC | Positive PCR Control |
| H12 | SA_00103 | PPC | Positive PCR Control |

**Table S3: List of cytokines and chemokines in Qiagen array used *in vitro* in Figure 2c*.***

| Data quality control (QC) | |
| --- | --- |
| Quality checks performed and results |  |
| Test Performed | **Test Result** |
| 1. PCR Array Reproducibility | All Samples Passed |
| 2. RT Efficiency | All Samples Passed |
| 3. Genomic DNA Contamination | All Samples Passed |

**Table S4:** Quality control of the Qiagen array used *in vitro* in Figure 2C*.*

| Normalization analysis | | | | | |
| --- | --- | --- | --- | --- | --- |
| Automatic selection from full panel | | | | | |
| Groups | Samples | Bmp4 | Cntf | Geometric Mean | Average Geometric Mean |
| Control Group | IFNg | 21.191742 | 26.935425 | 23.89 | 24.05 |
| Control Group | IFNg | 21.209953 | 26.528824 | 23.72 |  |
| Control Group | IFNg | 21.926346 | 27.47536 | 24.54 |  |
| Group 3 | HKMTI-1-005 | 21.146624 | 27.272264 | 24.01 | 23.87 |
| Group 3 | HKMTI-1-005 | 20.72649 | 26.189022 | 23.30 |  |
| Group 3 | HKMTI-1-005 | 21.67379 | 27.24161 | 24.30 |  |

**Table S5:** Normalisation analysis of the Qiagen array used *in vitro* in Figure 2c*.*

| **Patient No** | **Histology** | **Original stage** | **Disease status** |
| --- | --- | --- | --- |
| ASC19-026 | High-grade serous | 3B | Relapsed disease |
| ASC19-029 | High-grade serous | 3C | Primary |
| ASC19-030 | High-grade serous | 3C | Primary |
| ASC19-032 | High-grade serous | 3C | Primary |
| ASC19-033 | High-grade serous | 4B | Primary |
| ASC19-038 | High-grade endometrioid | n/a | Primary |
| ASC19-040 | High-grade serous | 3C | Primary |

**Table S6:** Patient clinical characteristics from Figure 2e results.

| Marker | | Fluorochrome | Dilution | Clone | Company | Code |
| --- | --- | --- | --- | --- | --- | --- |
| Myeloid | | | | | | |
| CD45 | Alexa Fluor 532 | | 1 in 100 | 30-F11 | ThermoFisher | 58-0451-82 |
| Ly6G | Alexa Fluor 700 | | 1 in 200 | 1A8 | Biolegend | 127622 |
| Ly6C | Brilliant Violet 605 | | 1 in 100 | HK1.4 | Biolegend | 128036 |
| MHC II | Brilliant Violet 510 | | 1 in 200 | M5/114.15.2 | Biolegend | 107636 |
| CD206 | Brilliant Violet 711 | | 1 in 100 | C068C2 | Biolegend | 141727 |
| CD80 | APC/Fire750 | | 1 in 100 | 16-10A1 | Biolegend | 104740 |
| CD86 | Brilliant Violet 785 | | 1 in 100 | GL-1 | Biolegend | 105043 |
| F4/80 | PE | | 1 in 50 | BM8 | Biolegend | 123110 |
| CD11b | eFluor450 | | 1 in 100 | M1/70 | ThermoFisher | 48-0112-82 |
| SiglecF | FITC | | 1 in 50 | REA798 | Miltenyi | 130-112-178 |
| CD11c | PerCP/Cy5.5 | | 1 in 100 | N418 | Biolegend | 117328 |
| PD-1 (CD279) | Brilliant Violet 605 | | 1 in 100 | 29F.1A12 | Biolegend | 135220 |
| Cxcr3 | PE/Dazzle 594 | | 1 in 100 | Cxcr3 - 173 | Biolegend | 126534 |
| Lymphoid | | | | | | |
| CD8a | Alexa Fluor 700 | | 1 in 400 | 53-6.7 | Biolegend | 100730 |
| CD19 | PerCP/Cy5.5 | | 1 in 200 | 6D5 | Biolegend | 115534 |
| CD4 | FITC | | 1 in 400 | RM4-5 | Biolegend | 100510 |
| CD44 | PE | | 1 in 200 | IM7 | Biolegend | 103024 |
| CD62L | Brilliant Violet 785 | | 1 in 100 | MEL-14 | Biolegend | 104440 |
| CD49b (DX5) | PE/Cy7 | | 1 in 100 | DX5 | Biolegend | 108916 |
| NK1.1 | APC | | 1 in 100 | PK136 | Biolegend | 108710 |
| Cxcr3 | Brilliant Violet 510 | | 1 in 100 | Cxcr3 - 173 | Biolegend | 126528 |
| PD-L1 (CD274) | Brilliant Violet 421 | | 1 in 100 | 10F.9G2 | Biolegend | 124315 |
| FOXP3 | Alexa Fluor 647 | | 1 in 50 | MF-14 | Biolegend | 126408 |

**Table S7:** List of conjugated antibodies used for immunophenotyping by flow cytometry in Figure 5 and Figure 6.

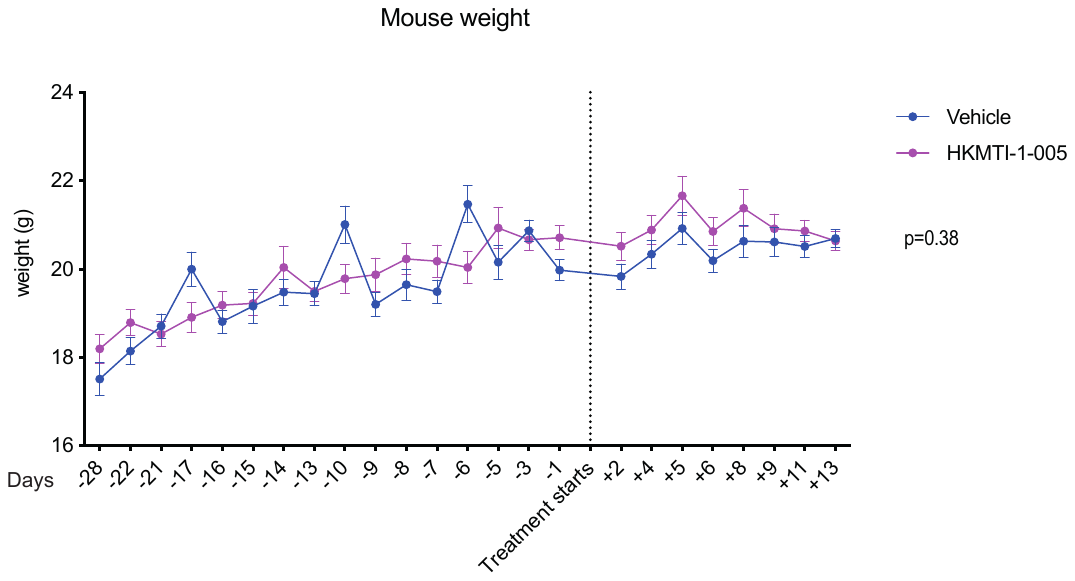

***Figure S4: Mouse weight during treatment***

Mouse body weight (n=24 per cohort) with either vehicle (1% Tween/3.6% DMSO in 0.9% NaCl IP bd) or HKMTI-1-005 (20mg/kg IP bd). Median weight was 19.8g ± 0.19 *vs* 20.1g ± 0.18, p=0.38, with vehicle and HKMTI-1-005, respectively. Unpaired t-test was used to compare means, bars show standard error of mean. IP: intraperitoneal, bd: twice daily.

***Figure S5: Immunohistochemistry of omental tumours harvested immediately after HKMTI-1-005 treatment***

Tumours harvested immediately after completion of HKMTI-1-005 treatment (as per Fig 4a) were stained for CD3, NKp46 (NK cell marker) and FoxP3 by immunohistochemistry. We observed an increase in both CD3 (H-score 14.8 ± 1.8 *vs* 20.4 ± 2.8, p=0.09) and NKp46 staining (3.4 ± 0.4 *vs* 4.6 ± 0.9, p=0.08) and a decrease FoxP3 staining (13.1 ± 2.64 *vs* 9.6 ± 2.07, p=0.3).

**Supplementary Methods**

*Immunohistochemistry*

Immunohistochemical (IHC) staining for CD3, NKp46 and Foxp3 was performed on 4 μm formalin fixed paraffin embedded (FFPE) sections that had previously been heated in an oven at 60C^o^ for 2 hours. IHC staining for CD3 was performed on a Leica Bond Rx Autostainer and staining for NKp46 and Foxp3 was performed on an Agilent Autostainer Link48. The antibodies were used at previously optimised dilutions: CD3 (Abcam ab16669) at 1/100, NKp46 (R&D Systems AF2225) at 1/200 and FoxP3 (Cell Signalling 12653) at 1/200 dilution.

The stained sections were then digitally captured on a Leica SCN400f slide scanner and image analysis was performed using HALO software (Indica Labs). Using the HALO software, firstly, non-malignant areas of tissue were manually excluded. Images were then analysed for the protein of interest using the CytoNuclear analysis tool v1.6. Images were then automatically scored using the histoscore method as described by Kirkegaard et al. This

algorithm is embedded in HALO software and after grading the staining intensity as negative (0), weakly stained (1), moderately stained (2) and strongly stained (3), it uses the formula below to calculate a histoscore value ranging between 0 and 300:

sum of (1 x % cells stained 1) + (2 x % stained 2) + (3 x % stained 3).

*ATAC sequencing filtration strategy*

For the analysis of ATACseq results, a filtration strategy was devised whereby only those peaks that were present in 2/5 HKMTI-1-005 samples and in 3/6 control samples were considered to truly represent areas of open chromatin. These were 74,424/184,605 (40%) in the control samples and 273,774/ 487,409 (56%). Subsequently, all the overlapping peaks between this group of peaks were removed and a remaining 218,718/273,774 peaks (80%) in the HKMTI-1-005 group were further interrogated for the presence of genes. 15,636 unique genes with any length of overlap with these 218,718 peaks were found and further examined for overlap with RNA sequencing differentially expressed genes (DEGs), as depicted in volcano plot in **Fig 3c**.

*ERV sequencing analysis strategy*

To remove low-expressing ERVs from our sequencing data, we applied a filtration threshold whereby an ERV needed a count of at least n=10 in one or more samples (in either the vehicle or HKMTI-1-005 group) in order to be included in our analysis. The total number of ERVs detected was 61,184 but only 2,781 and 2,465 passed the above filtration threshold in the control and treatment groups, respectively. 2,118 ERVs were common to both groups and they were subjected to differential expression analysis using the DESeq2 package. Out of these 2,118 ERVs, 51 were differentially expressed at the 5% FDR threshold.
